# m^3^C is a mitochondrial mRNA modification which promotes tumor progression

**DOI:** 10.1101/2025.01.15.633161

**Authors:** Audrey Penning, Kun Wang, Margot Célérier, Evelyne Collignon, Irina Primac, Louis Van der Linden, Martin Bizet, Emilie Calonne, Bouchra Hassabi, Céline Hubert, Pascale Putmans, Frédéric Murisier, Lionel Larbanoix, Younes Achouri, Jie Lan, Ivan Nemazanyy, Sophie Laurent, Patrick Jacquemin, Chengqi Yi, Rachel Deplus, François Fuks

## Abstract

Mitochondrial mRNA modifications are suggested to play a role in fine-tuning mitochondrial gene expression and function. However, the epitranscriptomic landscape of mitochondrial mRNA (mt-mRNA) remains poorly explored. Here, we uncover N3-methylcytosine (m^3^C) as a novel mt-mRNA that is catalyzed by METTL8, an enzyme previously known to modify mt-tRNA. Transcriptome-wide mapping reveals that METTL8-dependent m^3^C is enriched in mt-mRNAs encoding complex I subunits of the respiratory chain. Additionally, METTL8 is highly expressed in various cancers, notably in cervical cancer. METTL8 depletion impairs cell migration *in vitro* and reduces tumor growth in mouse xenografts. Finally, transcriptomic analyses further link METTL8 expression to oncogenic pathways, mitochondrial functions and complex I activity. Together, our results reveal a novel mitochondrial mRNA modification that promotes cancer progression.

## INTRODUCTION

Messenger RNA (mRNA) epitranscriptome has recently been under the spotlight. N6- methyladenosine (m^6^A), the most common and abundant internal mRNA modification identified to date, broadly affects mRNA metabolism. However, much remains to be explored in order to establish a comprehensive map of the mRNA modification landscape.

Similar to cytoplasmic RNA, mitochondrial-encoded RNA contains modified nucleotides. The best described RNA modifications occur on mt-tRNA and mt-rRNA, where they notably play a role in mitochondrial translation. Alongside mt-tRNA and mt-rRNA, RNA modifications in mitochondrial messenger RNA (mt-mRNA) have been recently reported, and they are thought to play a role in fine-tuning mitochondrial gene expression and functions. However, the epitranscriptomic landscape of mitochondrial mRNA remains elusive, with only two modifications currently mapped (i.e., pseudouridine (Ψ)^16^, and N1-methyladenosine (m^1^A)^17^). The extent of mitochondrial mRNA modification landscape and their putative role in mitochondria remain therefore unknown.

Several RNA modifications occur in cytoplasmic and mitochondrial tRNA, including N3- methylcytosine (m^3^C). METTL2A/B and METTL6 are responsible for tRNA^Thr^ and tRNA^Ser^ methylation, respectively. Together with their cofactor DARLD3, METTL2A/B install m^3^C on tRNA^Arg(UCU)/(CCU)^. The biological importance of these enzymes has been recently highlighted, with DALD3 mutations being involved in epileptic encephalopathy and METTL6 playing a role in stem cell self-renewal and tumor cell growth. METTL8 was initially proposed to catalyze m^3^C in cytoplasmic mRNA; however, no specific target has been identified.

The recent finding that METTL8 localizes to the mitochondria, where it modifies mt-tRNA^Ser/Thr^, has sparked renewed interest in the role of this enzyme. Of note, in relation to its role in regulating mitochondrial translation, METTL8 has been suggested to influence pancreatic cancer cell growth and to regulate cortical neural stem cells in both the embryonic mouse brain and human forebrain organoids.

Here, we show that METTL8, alongside its role in modifying mitochondrial tRNA, also catalyzes m^3^C on mitochondrial mRNA. Strikingly, METTL8 influences the protein levels of its m^3^C-modified targets, particularly the complex I subunits of the respiratory chain, thereby affecting its activity. METTL8 is overexpressed in various cancers and correlates with poor clinical outcomes in cervical cancer. Finally, we show that METTL8-dependent m^3^C plays a role in cervical cancer growth in xenografts. More broadly METTL8-driven transcriptomic changes are enriched for oncogenic pathways, and mitochondrial functions. Overall, our results reveal a novel mt-mRNA modification (m^3^C) and its writing enzyme (METTL8), that play a role in mitochondria and cancer.

## RESULTS

### Mitochondrial mRNA is m^3^C-modified by METTL8

To get a broad view of the presence of m^3^C in different types of RNA encoded in the nucleus and in the mitochondria, we initially performed specific RNA enrichments to isolate total and poly(A) RNA from the mitochondrial and non-mitochondrial cellular fractions (Figure 1A). This fractionation strategy is key since poly(A) enrichment is usually performed on whole-cell RNA and therefore does not allow the distinction between RNA modifications from nuclear RNA and mitochondrial RNA. Proper enrichment of each RNA fractions has been confirmed by RT-qPCR (Figures 1B and S1A). We next assessed the global level of m^3^C in each fractions by mass spectrometry. As previously reported, m^3^C is found in both whole-cell total RNA (which includes the tRNA fraction known to be m^3^C modified), and in whole-cell poly(A) enriched RNA (Figure 1C)^1^. Since the whole-cell poly(A) fraction encompasses mRNA from both the nucleus and the mitochondria, we further unraveled which mRNA fractions is modified. Importantly, m^3^C signal was much higher in poly(A) RNA from the mitochondrial fraction (0,017% m^3^C/C vs 0,0004% m^3^C/C for the non-mitochondrial fraction). Therefore, the m^3^C signal observed in whole-cell poly(A) RNA, mostly, if not entirely, derives from mitochondrial mRNA (Figure 1C, right). To further confirm that m^3^C is essentially modifying mitochondrial mRNA, we generated Rho0 cells through ethidium bromide treatment^2^. Rho0 cells are devoid of mitochondrial DNA and consequently of mitochondrial RNA, as validated by qPCR (Figures 1D and S1B). In these cells, m^3^C levels were almost abolished in poly(A) RNA, with only 0.00006% of m^3^C/C remaining (Figure 1E). These results further suggest that the presence of m^3^C in whole-cell poly(A) RNA derives from the modified mitochondria-encoded mRNAs and not from the nuclear-encoded mRNAs.

**Figure 1.**
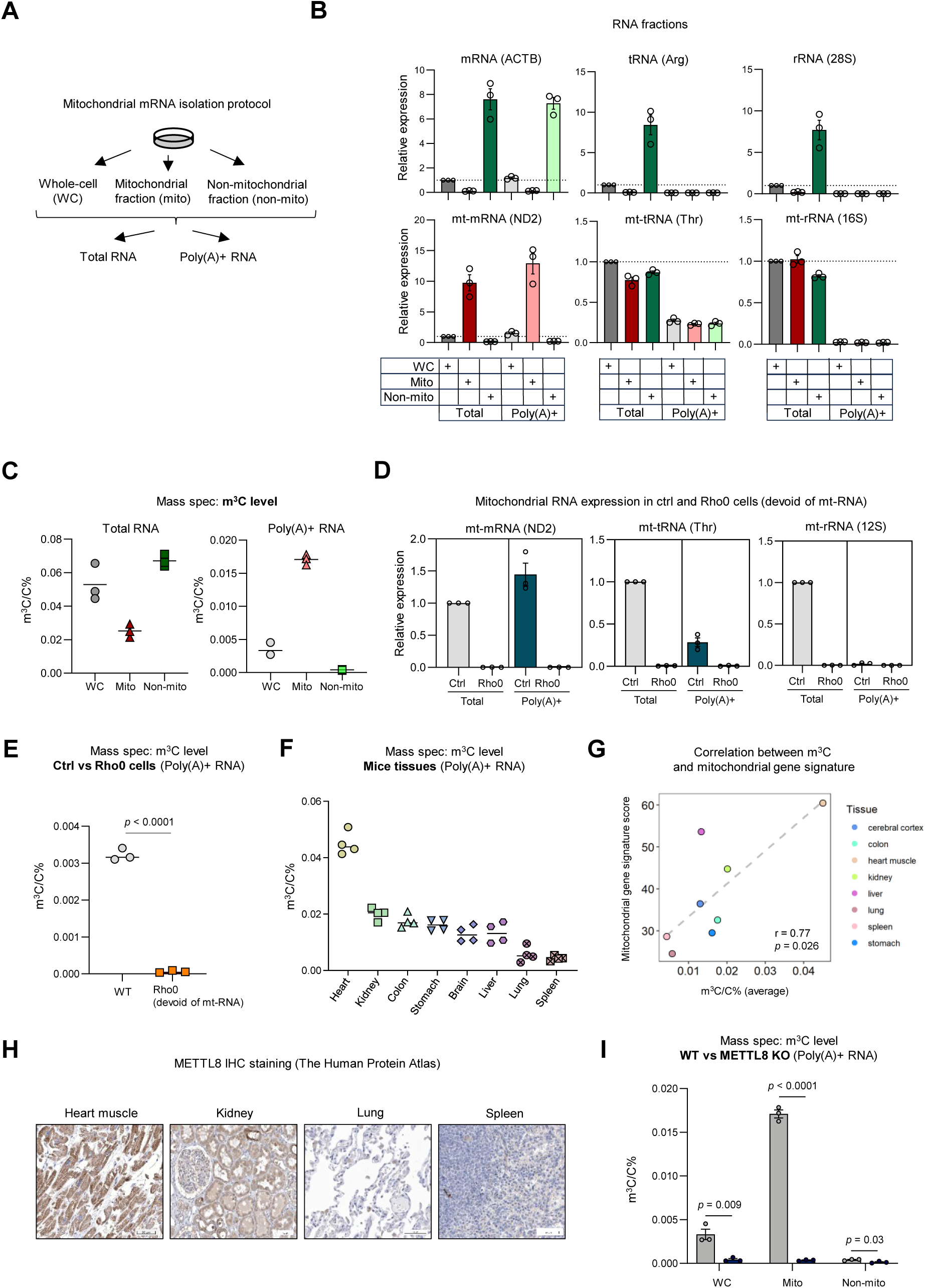
m^3^C modification in mitochondrial mRNA. (A) Scheme illustrating the protocol of total and poly(A) RNA enrichment from whole-cell, mitochondrial and non-mitochondrial fractions. (B) Relative quantification of mRNA (ACTB), tRNA (tRNA Arg), rRNA (28S), mt-mRNA (ND2), mt-tRNA (mt-tRNA Thr) and mt-rRNA (16S) transcripts from the total RNA and poly(A) RNA from whole-cell (WC), mitochondrial and non-mitochondrial fractions. (n=3, mean± SEM) (C) Quantification of m^3^C (m^3^C/C%) in total and poly(A) RNA from whole-cell (WC), mitochondrial and non-mitochondrial fractions. (*n*=3, mean± SEM) (D) Relative quantification of mt-mRNA (ND2), mt-tRNA (mt-tRNA Thr), and mt-rRNA (12S) transcripts from control and Rho0 cells, in total RNA and poly(A) RNA. (n=3, mean± SEM) (E) Quantification of m^3^C (m^3^C/C%) in poly(A) RNA from WT and Rho0 cells. (*n*=3, mean± SEM, two-tailed *t* test) (F) m^3^C level (m^3^C/C%) in poly(A) RNA from a panel of mice tissues (heart, kidney, colon, stomach, brain, liver, lung, spleen). (*n*=4 mice, mean±SEM) (G) Correlation between m^3^C level and mitochondrial genes signature score in various mouse tissues (cerebral cortex, colon, heart muscle, kidney, liver, lung, spleen, stomach). Pearson correlation coefficient (r) and *p*-value as indicated. (H) Quantification of m^3^C in poly(A) RNA from whole-cell, mitochondrial and non-mitochondrial fractions in WT and METTL8 KO cells. (*n*=3, mean± SEM, two-tailed *t* test) (I) Immunohistochemistry (IHC) staining of METTL8 in heart muscle, kidney, lung and spleen (image credit: Human Protein Atlas).

To evaluate the presence of this mark on mt-mRNA *in vivo*, we measured the global level of m^3^C in poly(A) enriched RNA from various mouse tissues. As shown in Figure 1F, m^3^C modifies poly(A) enriched RNA from all tissues, with the highest level observed in tissues known to be enriched in mitochondria (e.g., heart)^3^. To further validate this observation, we assessed mitochondrial content across mouse tissues using a gene signature for mitochondrial genes. Using this method, we found a significant correlation between m^3^C level and mitochondrial signal among these tissues (Figure 1G). This further suggests that m^3^C level is associated with mitochondrial content, due to its presence in mitochondrial RNA.

METTL8 was recently shown to be located in mitochondria and to catalyze m^3^C in mitochondrial RNA^4^. We used the Human Protein Atlas and observed that METTL8 protein staining is high in tissues with abundant mitochondrial content, such as the heart and kidney, and low in tissues with lower mitochondrial content, like the lung and spleen (Figures 1F-1H). Since METTL8 catalyzes m^3^C in mt-tRNAs^4^, we next assessed if this enzyme could be the m^3^C methylase of mitochondrial mRNA. Therefore, we generated HeLa METTL8 knockout cells by CRISPR/Cas9. We validated the loss of METTL8 by western blot in all cellular fractions (whole-cell, mitochondrial and non-mitochondrial), with TOMM20 and ACTB being used as controls for mitochondrial and non-mitochondrial enrichment, respectively (Figure S1C). Of note, METTL8 was localized in the mitochondria in WT cells, as reported^4^. Next, we assessed the level of m^3^C in poly(A) enriched RNA from whole-cell, mitochondrial and non-mitochondrial fractions in WT and METTL8 KO cells. As shown in Figure 1I, m^3^C is almost completely abolished in all three fractions upon METTL8 KO, with the most drastic difference being observed in mitochondria, where m^3^C is the most abundant (0,017% m^3^C/C vs 0,00036% m^3^C/C). The observed reduction of m^3^C signal in poly(A) RNA from non-mitochondria fractions is likely due to the METTL8-dependent m^3^C-modified mt-tRNAs, that remain after the poly(A) enrichment (as previously measured by RT-qPCR, see Figures 1B and S1A). Together, these results indicate for the first time that m^3^C is present in mitochondrial mRNA and depends on METTL8.

### METTL8 is overexpressed in various cancers and correlates with proliferative and mitochondrial gene signatures

It has been well established that mitochondria are important mediators of various tumorigenic mechanisms^5^. This, and the previously reported roles of mitochondrial METTL8 in pancreatic cancers growth *in vitro*^4^, led us to speculate that METTL8 might play a role in various cancer types. First, to get a broad view of potential dysregulations of METTL8 in cancers, we investigated METTL8 expression in tumor samples and their healthy counterparts using The Cancer Genome Atlas (TCGA) database. We found a higher expression of METTL8 in several cancers (cervical, cholangiocarcinoma, lung) (Figure 2A). This overexpression was also reflected by METTL8 protein staining in cervical and lung cancers (The Human Protein Atlas) (Figure 2B). To further substantiate these observations, we quantified the expression of METTL8 by RT-qPCR, in a cohort of human cervical cancer biopsies (20 healthy and 19 cancer tissues). As shown in Figure 2C, METTL8 expression is higher in tumor biopsies compared to healthy cervical tissues. Additionally, high METTL8 expression was associated with poor survival in public datasets of several cancers (cervical, lung, ovarian and liver) (Figures 2D and S2A-S2C).

**Figure 2.**
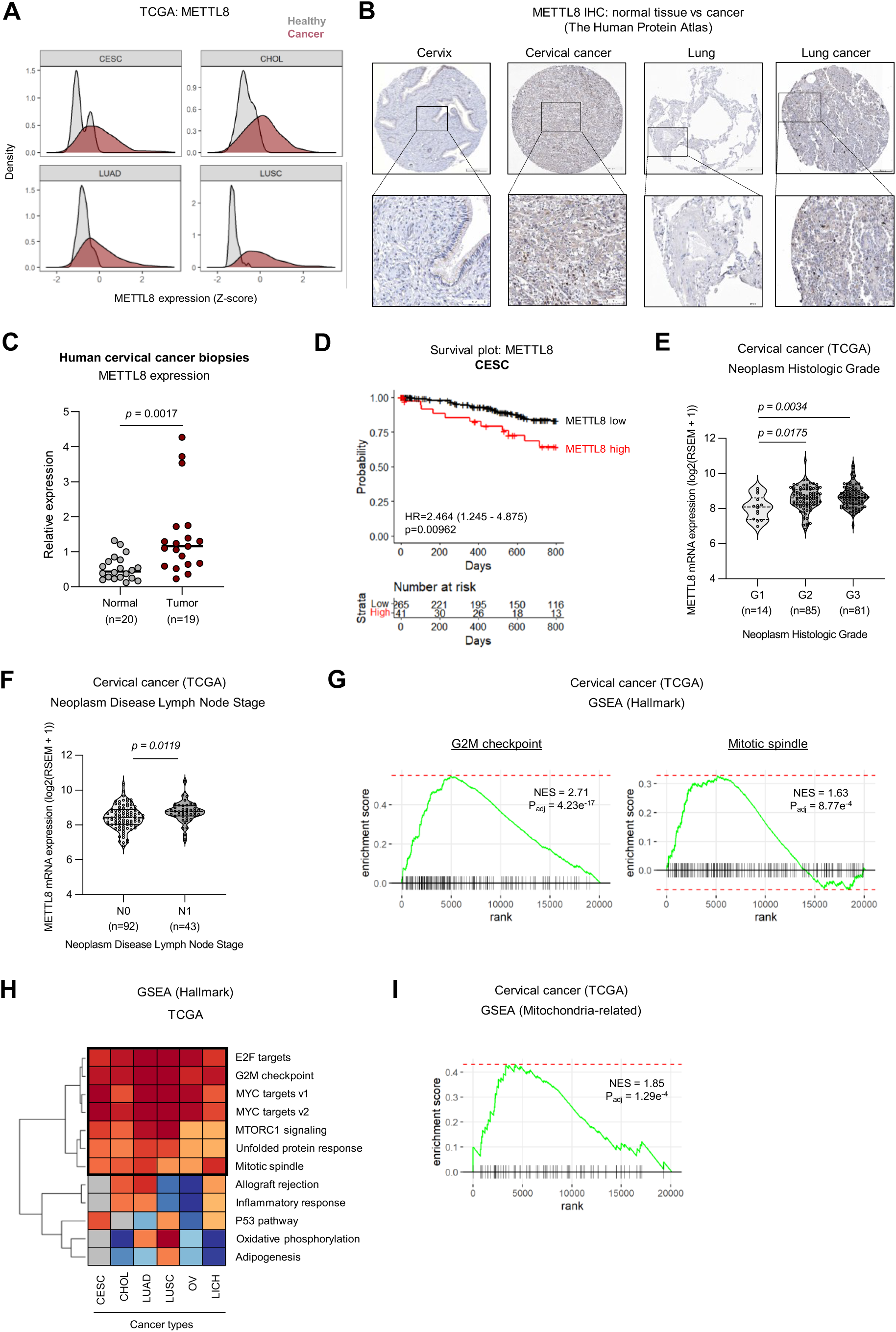
Overexpression of METTL8 in cancers. (A) Density plot of METTL8 expression (Z-score) in various healthy and cancer tissues from the TCGA cohort (The Cancer Genome Atlas) (CESC, CHOL, LUAD and LUSC). (B) IHC staining of METTL8 in healthy cervix versus cervical cancer tissues; and healthy lung versus lung cancer tissues (image credit: Human Protein Atlas). (C) Relative quantification of METTL8 expression in human cervical cancer biopsies and normal tissues. The number of samples (*n*) is indicated for each group. (Two-tailed *t* test) (D) Kaplan-Meier curves showing the overall survival of cervical cancer patients categorized as METTL8 high or low. (Hazard Ratio (HR) with 95% confidence interval and logrank test) (E) METTL8 expression in cervical cancer neoplasm categorized by histologic grade in TCGA. The number of samples (*n*) is indicated for each group. (One-way ANOVA with Tukey’s multiple comparisons test, violin plots showing all replicates) (F) METTL8 expression in cervical cancer neoplasm disease lymph node stage in TCGA. The number of samples (*n*) is indicated for each group. (Two-tailed *t* test, violin plots showing all replicates) (G-H) GSEA in TCGA cancers using gene sets of the ‘Hallmark’ collection. Genes were pre-ranked based on correlation with METTL8 expression. Examples are shown for cervical cancer (G). Heatmap illustrating the NES (normalized enrichment score) in several cancer types (CESC, CHOL, LUAD, LUSC, OV, LICH) (H). (I) GSEA in TCGA cervical cancer using a signature of mitochondria-related genes. All genes were pre-ranked based on correlation with METTL8 expression.

Prompted by our findings in cervical cancers (TCGA) and the role of METTL8-mediated m^3^C in HeLa cells (Figure 1I), we further investigated the impact of METTL8 in cervical cancer progression. Interestingly, we observed a significant correlation between METTL8 expression and increased histological grade (G1 to G3) (Figure 2E), as well as with lymph nodes status (N1 vs N0) (Figure 2F) in TCGA. To expose the mechanisms through which METTL8 might favor cervical cancer progression, we applied gene set enrichment analysis (GSEA) to TCGA data and identified ‘G2M checkpoint’ and ‘Mitotic spindle’ as enriched gene sets positively correlating with METTL8 expression (Figure 2G), suggesting that METTL8 may promote oncogenic pathways.

To further extend these findings beyond cervical cancer, we performed GSEA analysis on several publicly available cancer data sets from TCGA (cholangiocarcinoma, lung, ovarian, liver). We identified several key oncogenic pathways, including ‘E2F targets’, ‘G2M checkpoint’, ‘MYC targets’, and ‘MTORC1 signaling’ which were consistently overrepresented in association with METTL8 expression across these cancer types (Figure 2H).

Finally, given the role of METTL8 as the m^3^C writer in mitochondrial RNA, we identified by GSEA in several cancer types a positive correlation between METTL8 expression and a mitochondria-related gene signature (Figure 2I and S2D-S2E).

Overall, we found that METTL8 is overexpressed in various cancers, including in cervical cancer, where high expression is associated with disease progression. Furthermore, various key oncogenic gene sets and mitochondrial signal both correlate with METTL8 expression in those cancers.

### METTL8 catalyzes m^3^C on mitochondrial mRNA from complex I

Having found that m^3^C is a novel modification present in mt-mRNA, we next investigated which mitochondrial mRNA are modified in cervical cancer HeLa cells. Therefore, we performed m^3^C-RIP-seq: immunoprecipitation of m^3^C-containing mRNA with an anti-m^3^C antibody followed by next-generation sequencing.

First, we validated our approach by performing m^3^C-RIP-qPCR on WT cells, showing the specific enrichment of known m^3^C modified tRNAs versus unmodified (Figure 3A). We then performed m^3^C- RIP-seq in WT and METTL8 KO cells (Figure 3B), by focusing our analysis on the mt-mRNA, since this fraction showed the highest reduction in m^3^C upon METTL8 knockout (Figure 1I). This approach identified 7 mt-mRNAs as being m^3^C-modified among the 13 mRNAs encoded by the mitochondria (log_2_ fold change < -0.8; *p*-value < 0.05). Strikingly, 6 out of these 7 are subunits of the complex I: ND1, ND2, ND4, ND4L, ND5 and ND6 (Figure 3C). Examples of m^3^C enrichment profiles in WT and METTL8 KO are shown in Figure 3D, with negative examples (mt-mRNAs not m^3^C modified) being shown in Figure S3A. Lastly, since mt-mRNAs are endogenously produced from a polycistronic precursor mitochondrial RNA, we sought to exclude the possibility that the observed m^3^C peaks resulted from incomplete cleavage of the well-known m^3^C-modified mt-tRNAs. Therefore, we checked whether these modified mt-tRNAs were in close proximity to our identified mt-mRNA targets. As represented in Figure 3E, our mt-mRNA targets are not found in proximity of known mt-tRNA targets, suggesting that the m^3^C peaks we observed do not result from enrichment of the precursor RNA. Overall, we established the first map of m^3^C in mt-mRNA and revealed 7 mt-mRNAs as m^3^C modified by METTL8.

**Figure 3.**
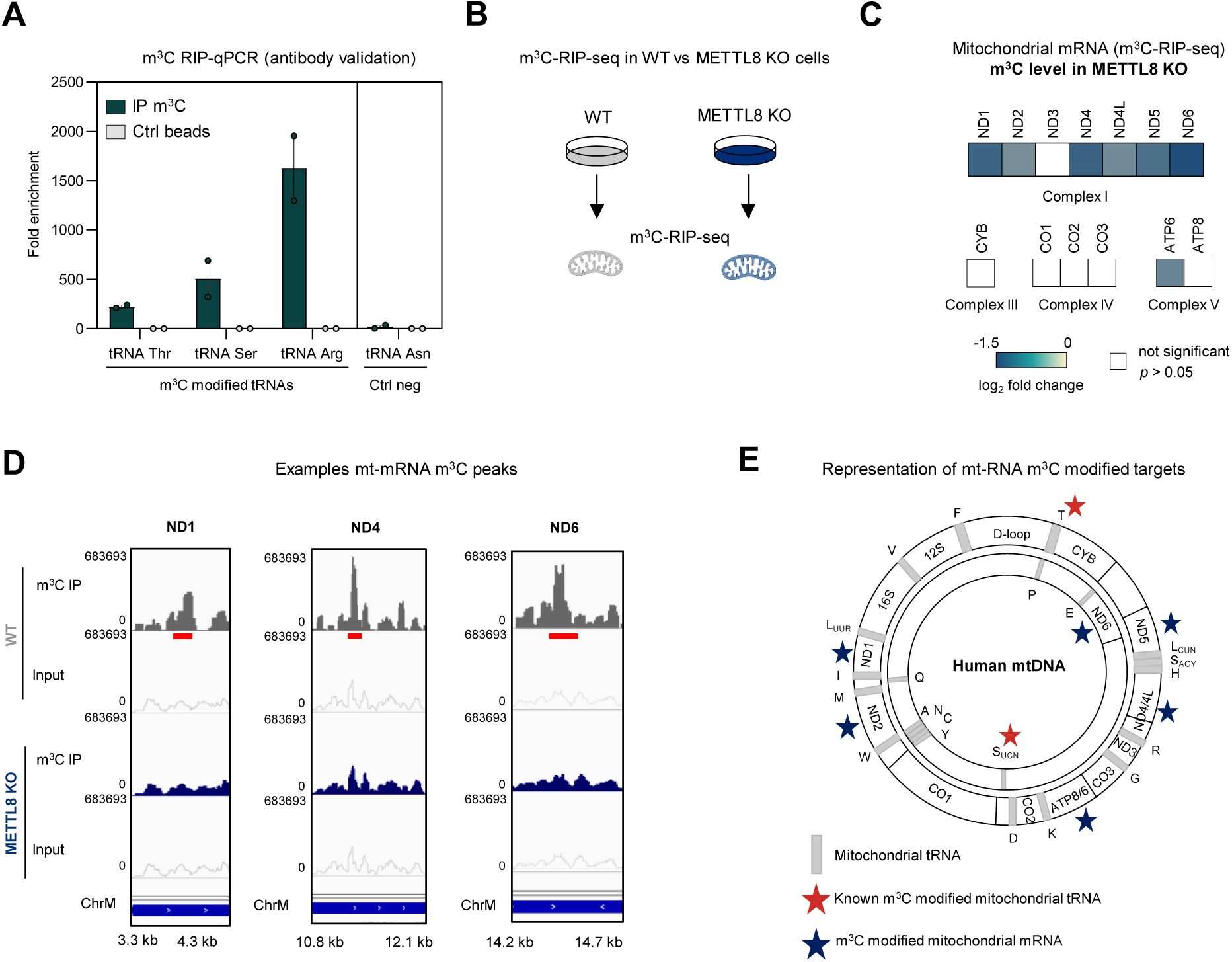
m^3^C landscape in mt-mRNA. (A) m^3^C-RIP-qPCR on positive m^3^C modified tRNAs (Thr, Ser and Arg) and a negative unmodified control tRNA (Asn). (*n*=2, mean±SEM) (B) Scheme illustrating the m^3^C-RIP-seq experimental design using WT and METTL8 KO cells. (C) Reduced m^3^C peaks level in METTL8 KO in mt-mRNA. (*p*-value corrected for multi-testing < 0.05) (D) Examples of m^3^C-RIP-seq tracks (ND1, ND4 and ND6) in WT and METTL8 KO cells. Red lines indicate the peak locations. (E) Scheme representing the mitochondrial genome with genes coding for the previously described m^3^C modified mt-tRNAs labeled with red stars and genes coding for the newly identified m^3^C modified mt-mRNAs labeled with blue stars.

### METTL8-dependent m^3^C modulates complex I activity

To identify the potential function of m^3^C, we first assessed the expression of all mt-mRNAs by RT-qPCR (Figure S4A). However, we did not find any difference between WT and METTL8 KO, suggesting that m^3^C does not affect mitochondrial gene expression at the RNA level. We next assessed the protein level of m^3^C targets by western blot. Depletion of m^3^C upon METTL8 knockout resulted in a higher level of ND1 and lower level of ND4 and ND6 (Figures 4A-4C), with no effect on an unmodified negative control (CO2, see Figure S4B). This suggests that m^3^C can have varied effects on its targets.

**Figure 4.**
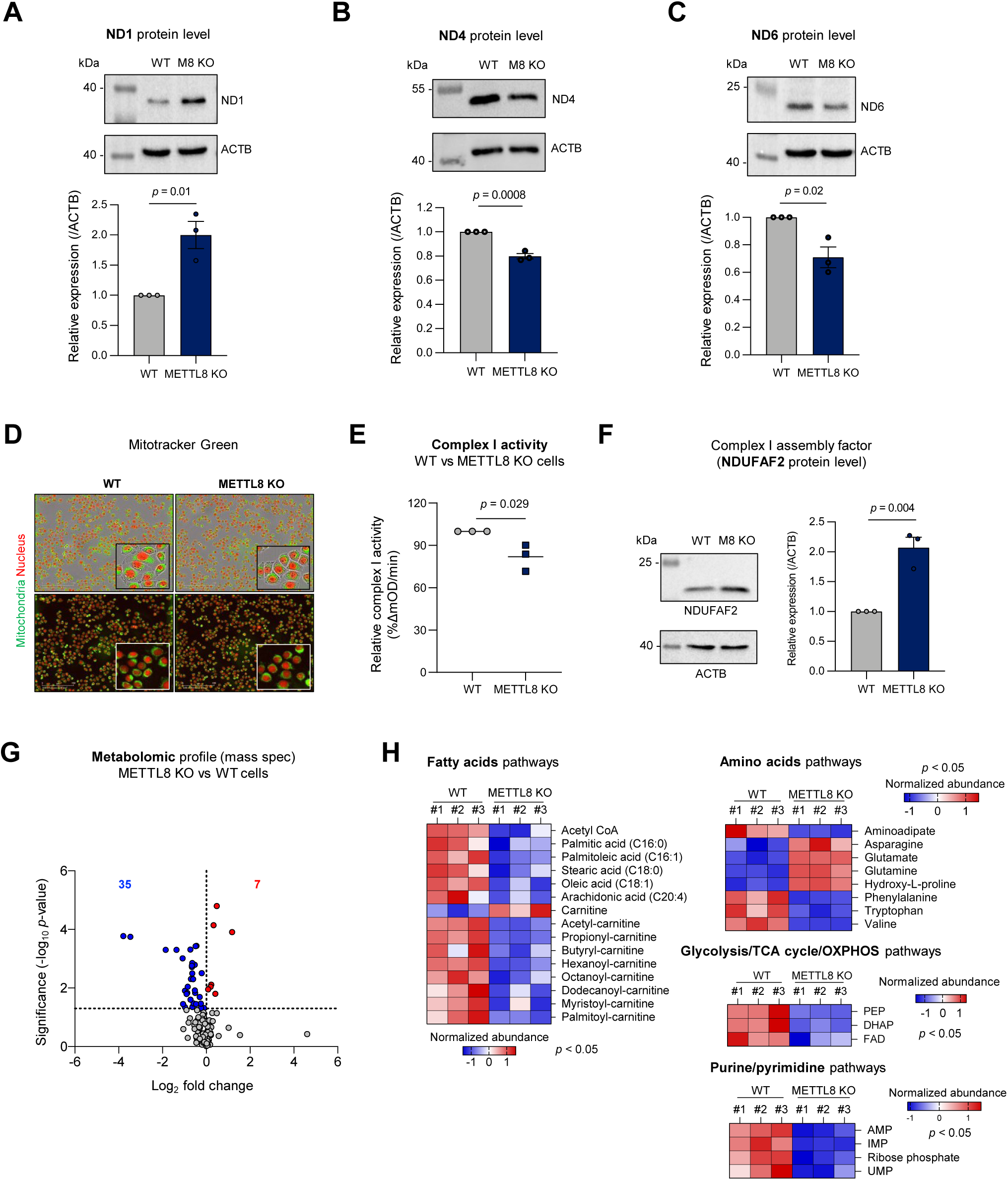
Functions of METTL8-dependent m^3^C. (A-C) Representative western blot images of ND1 (A), ND4 (B) and ND6 (C) in WT and METTL8 (M8) KO HeLa cells with protein quantification. (level normalized to ACTB, *n*=3, two-tailed *t* test, mean±SEM) (D) Mitotracker Green signal (mitochondrial staining) and Nuclight Red signal (nuclear staining) in WT and METTL8 KO cells. Representative pictures are shown (*n*=3) (E) Relative activity of complex I (colorimetric microplate assay, %ΔmOD/min, see Methods) in WT and METTL8 KO HeLa cells. (*n*=3, two-tailed *t* test, mean±SEM) (F) Representative western blot images of NDUFAF2 in WT and METTL8 (M8) KO HeLa cells with protein quantification. (normalized to ACTB, *n*=3, two-tailed *t* test, mean±SEM) (G) Volcano plot showing the differential metabolites upon METTL8 depletion. (*n*=3, two-tailed *t* test, *p*-value < 0.05) (H) Examples of significantly dysregulated metabolites in METTL8 depleted cells, classified in ‘fatty acids’, ‘amino acids’, ‘glycolysis/TCA cycle/OXPHOS’ and ‘purine/pyrimidine’ pathways. (*n*=3, two-tailed *t* test, *p*-value < 0.05)

Next, we explored the global effect of METTL8-dependent m^3^C on mitochondria. First, we assessed mitochondrial content using the mitotracker dye (Figure 4D) and found no change upon METTL8 depletion (Figure S4C). We confirmed this finding by mt-DNA quantification (Figure S4D).

Since most m^3^C-modified mt-mRNAs encode subunits of the complex I of the electron transport chain (Figure 3C), we next investigated whether METTL8 specifically regulates the complex I. As shown in Figure 4E, METTL8 depletion leads to reduced activity of this complex *in vitro*. Furthermore, expression of the assembly factor NDUFAF2, known to be upregulated in response to complex I deficiency^6^, is higher in METTL8 KO cells compared to WT, further suggesting a deficiency in complex I upon METTL8 depletion (Figure 4F).

Considering the well-known and important role of mitochondria in metabolism^7^, we next performed metabolomics profiling in WT and METTL8 KO cells (Figure 4G). We observed numerous alterations of different amino acids and acylcarnitine levels upon METTL8 knockout (Figure 4H), as previously described in a complex I deficient mouse model (Ndufs4-/-)^8^. Specifically, we observed an important decrease of fatty acid acylcarnitine of even and odd chain length, as well as 2- aminoadipate (Figure 4H). The observed decreased level of these metabolites may result from increased electron flux through the mitochondrial flavoprotein dehydrogenases to the electron transfer flavoprotein (ETF/ETF-QO) system, and *in fine* to the Q pool of the respiratory chain, compensating for the reduced complex I electron transfer, as previously suggested^8^. Overall, we observed a dysregulation in various metabolites, mostly reflecting fatty acids catabolism, potentially as a compensatory mechanism to complex I deficiency.

Taken together, our findings indicate that METTL8-dependent m^3^C affects the protein level of its targets and regulates mitochondrial complex I activity, as notably evidenced by the altered metabolome upon METTL8 depletion.

### METTL8 promotes cervical cancer

Considering the emerging crosstalk between cervical cancer and mitochondria^9^, along with the high levels of METTL8 in this cancer type, we aimed to investigate for the first time the potential pathological role of m^3^C in cervical cancer.

We first investigated how METTL8 influences HeLa cell migration. Using the Incucyte Scratch Wound cell migration assay, we found that METTL8 depleted cells display lower ability to migrate (Figure 5A). Furthermore, we confirmed that the effect observed in migration was not due to a bias in proliferation rate, since we did not observe any substantial difference in proliferation (Figure S5A). To investigate the role of METTL8 in tumor growth *in vivo*, we injected WT and METTL8 KO HeLa cells in the flanks of immunodeficient mice and tracked tumors sizes. As shown in Figure 5B, METTL8 depletion markedly reduced tumor growth, suggesting that METTL8-dependent m^3^C could play a role in tumorigenesis. We further confirmed the role of METTL8 in promoting cervical cancer growth *in vivo* using luciferase-labeled xenografts (Figure 5C).

**Figure 5.**
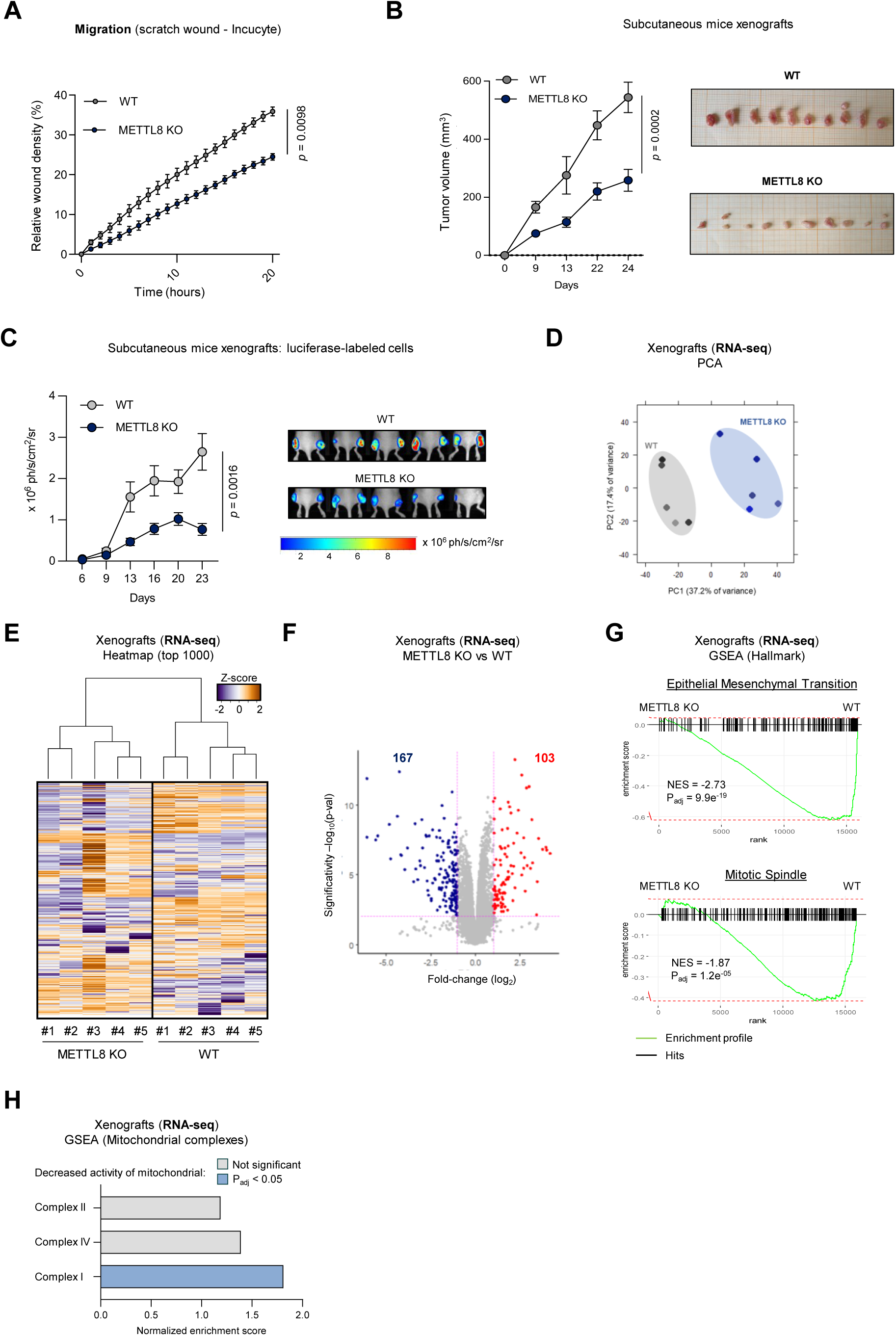
Pathological role of METTL8 in cervical cancer. (A) Real-time migration of WT and METTL8 KO cells. (*n*=3, two-way ANOVA, mean±SEM) (B) Tumor volume of WT and METTL8 KO cervical cancer xenografts. (*n*=10 tumors per condition, two-way ANOVA, mean±SEM) (C) Bioluminescent imaging of luciferase-labeled WT and METTL8 KO HeLa xenografts. (*n*=10 tumors per condition, two-way ANOVA, mean±SEM) (D) PCA plot for RNA-seq in xenografts, showing that WT and METTL8 KO conditions have distinct transcriptomic profiles. (*n*=5 tumors per condition) (E) Heatmap of the top 1000 most variable genes by RNA-seq in xenografts, with hierarchical clustering segregating WT and METTL8 KO tumors. (*n*=5 tumors per condition) (F) Volcano plot showing genes differentially expressed upon METTL8 depletion by RNA-seq. (|Log_2_ fold change| > 1 and adjusted *p*-value < 0.05) (G-H) GSEA in xenograft RNA-seq data. Genes were pre-ranked based on differential expression (fold change) upon METTL8 depletion. Examples of gene sets from the ‘Hallmark’ collection (G) and mitochondrial signatures of ‘decreased activity’ of complexes I, II and IV (H) are shown.

To identify dysregulated pathways driving the observed phenotype, we performed RNA-seq in xenografts tumors. Overall, METTL8 KO tumors display a distinct transcriptome profile (Figures 5D-5E). We identified 270 transcripts significantly differentially expressed upon METTL8 depletion, with both gain and loss of expression (Figure 5F). We then applied GSEA and identified ‘Epithelial Mesenchymal Transition’ and ‘Mitotic Spindle’ as two of the top underrepresented gene sets in METTL8 depleted cells (Figure 5G). These results suggest that METTL8 promotes pathways critical for sustaining tumorigenesis. Interestingly, GSEA also uncovered a link between METTL8 and its mitochondrial role in cervical cancer xenografts, since ‘abnormal activity of mitochondrial respiratory chain’ and ‘decreased activity of mitochondrial complex I’ were both overrepresented gene sets in METTL8 depleted cells (Figures S5B-S5C). Consistent with our previous observation that METTL8-mediated m^3^C targets in mt-mRNA are almost exclusively from the complex I (Figure 3C), we observed a significant association between METTL8 depletion and ‘decreased activity of complex I’, whereas no such correlation was found for complex II or IV (Figure 5H).

Overall, our results suggest that METTL8 plays an oncogenic role in cervical cancer through its regulation of m^3^C modified mt-mRNA metabolism and, in particular, complex I activity, thereby supporting mitochondrial respiration.

## DISCUSSION

A new area of epigenetic research is on the rise: mRNA epitranscriptomics. This field has increasingly intrigued researchers due to the numerous existing modifications identified in tRNA or rRNA, and recently reported in mRNA, as well as their involvement in the regulation of major biological processes^10–15^. The vast majority of mRNA epitranscriptomic studies have focused on cytoplasmic mRNA, leaving the mitochondrial mRNA modification landscape largely unexplored. To date, only two modifications have been reported in mitochondrial mRNA: pseudouridine (Ψ)^16^ and N1-methyladenosine (m^1^A)^17^. The extent of mitochondrial mRNA modifications and their putative role remain therefore largely unknown. Considering the crucial role of mitochondria in a plethora of cellular pathways, we sought to investigate the poorly characterized mitochondrial mRNA modification landscape.

In 2017, Xu *et al.* reported that m^3^C exists on poly(A) RNA and is catalyzed by the methylase METTL8^1^. Two subsequent studies showed that METTL8 is located in the mitochondria where it catalyzes m^3^C on two tRNAs^4,18^. Of note, Xu and colleagues used whole cells poly(A) RNA enrichment and global mass spectrometry to assess the presence of m^3^C^1^. Yet, whole cells poly(A) RNA enrichment comprises nuclear-encoded poly(A) RNA as well as mitochondrial-encoded poly(A) RNA. The latter is unfortunately frequently omitted in epitranscriptomic studies relying solely on global mass spectrometry. Therefore, we first conducted an in-depth assessment of the presence of m^3^C in various RNA species transcribed in the nucleus and in the mitochondria. We found that, while this mark is indeed present in poly(A) RNA, it is most abundant in the mitochondria.

We further extended our study of m^3^C by unravelling its mitochondrial mRNA landscape. Previous transcriptome-wide mapping studies did not detect m^3^C in mRNA, and suggested that this might be due to the substoichiometric level of m^3^C, present only in a small subset of mRNAs^19,20^. In contrast to these studies that relied on chemical mapping, we used m^3^C-RIP-seq, a method involving an enrichment step with a m^3^C antibody that could facilitate the observation of this lowly abundant modification. Alongside this enrichment step, we applied two important validation steps to our transcriptome-wide analysis: (1) we validated our antibody using known m^3^C modified tRNAs; and (2) we only considered mitochondrial mRNAs with reduced m^3^C level upon METTL8 depletion.

Overall, we found 7 mitochondrial mRNA targets that appear to be m^3^C modified, further confirming that it is restricted to a small number of mRNAs. This could explain why the mark has not been found with previous total RNA assessment. Therefore, it would be of interest to perform chemical mapping methods after specific enrichment of mitochondrial poly(A) RNA to overcome the substoichiometric level of m^3^C on mRNA.

Mechanistically, after having found that m^3^C is present in seven mitochondrial mRNAs, our data evidence that m^3^C might affect the protein level of its targets. In line with previous reports on other modifications affecting Watson-Crick base pairing (e.g., m^1^A)^17^, the depletion of m^3^C upon METTL8 KO results in a higher level of ND1 protein. However, it appears that the consequence of m^3^C varies, since METTL8 depletion has the opposite effect on ND4 and ND6, for example. An association between METTL8-dependent m^3^C in mt-tRNAs and mitochondrial translation has been also recently reported^4^, raising another level of complexity in the regulation of protein level. If the effects of METTL8 depletion on ND6 is similar between studies; there is an opposite regulation of ND1. This difference could be explained by the dual role of METTL8 as the m^3^C writer on mt-mRNAs but also on mt-tRNAs, which could be more prominent according to the different contexts. Indeed, our data were obtained in HeLa cells, when Schöller and colleagues used 293T cells^4^, possibly explaining the differences in METTL8-dependent m^3^C functions. The authors also found functional differences between two pancreatic cancer cell lines, showing that the role of METTL8 is probably cell type-dependent^4^. Furthermore, Zhang et al. recently addressed m^3^C and METTL8 in a neurological context and found CO2 and ATP6 to be dysregulated upon METTL8 depletion, which were not pinpoint by previous studies. This discrepancy further demonstrates the importance of context in assessing the role of mRNA modifications^21^. Finally, m^3^C could potentially also exerts its role through the recruitment of various readers with different functional consequences of the modification, as has been described for m^6^A^14^. However, further studies are warranted to unravel this hypothesis. Despite the functional differences between studies, one major common observation is that METTL8-dependent m^3^C has an impact on mitochondrial translation. Functionally, METTL8-dependent m^3^C seems to affect mitochondrial complex I activity and various metabolic pathways such as the acylcarnitine pathways. This corroborates that m^3^C is most present on mt-mRNA from complex I (6 out of 7 targets).

Altogether our data suggest that the epitranscriptomic field goes beyond the cytoplasmic mRNAs and that mitochondrial mRNAs can also be modified, thereby affecting key mitochondrial processes. Our results open the possibility of identifying additional modifications on mitochondrial mRNA, which could play a crucial role in mitochondrial metabolism.

mRNA modifications have increasingly been linked to cancer pathogenesis^15,22^. A handful of recent reports describe the contribution of mitochondrial RNA modifications to cancer. Among these, several studies report a potential implication of METTL8 in regulating colorectal^23^ and pancreatic^4^ cancer cells proliferation *in vitro*. However, the role of METTL8 and m^3^C in cancer remains poorly explored. Furthermore, while a mitochondrial tRNA modification 5-methylcytosine (m^5^C) has recently been highlighted as a key actor in tumor malignancy by promoting metastasis^24^, the role of mitochondrial mRNA modifications in disease remains a mystery.

Strikingly, METTL8 is overexpressed in several tumor types, thus we decided to tackle the putative pathological role of m^3^C in cancer. Given the emerging crosstalk between cervical cancer and mitochondria^9^, and the role of METTL8-dependent m^3^C in HeLa cells (Figures 1, 3 and 4), we focused on cervical cancer. Overall, our data show that METTL8-dependent m^3^C promotes cervical cancer. Interestingly, alongside its role in mitochondrial neuro-muscular disorders, complex I impairment is a major contributing factor in pathological processes, including cancer. It has notably been shown that compromising complex I activity might impaired tumorigenesis^25–27^. However, a link with mitochondrial mRNA modifications has never been established. Therefore, our study, alongside the recently described role of m^5^C in mitochondrial tRNA^24^, opens the prospect of a novel regulatory layer of mitochondrial epitranscriptomics in tumorigenesis.

Altogether our findings document for the first time the presence and the role of a mitochondrial mRNA modification. We pave the way for the discovery of additional modifications decorating mitochondrial mRNA, which could play major roles in cell biology. Overall, our findings sheds a new light on the role of mitochondrial mRNA epitranscriptomics in cancer, with mechanistic and therapeutic potentials, and thereby open the frontiers of the epitranscriptomic field beyond cytoplasmic mRNA modifications.

## ACKNOWLEDGMENTS

A.P., M.C., E. Collignon, I.P., L.V.D.L., M.B., B.H., C.H., and F.M. were supported by the Belgian FRS-FNRS, FRIA, or Télévie. F.F. is a ULB Professor. R.D. is a ULB lecturer. The CMMI is supported by the European Regional Development Fund and the Walloon Region. F.F.’s lab was funded by grants from the FNRS and Télévie, the “Action de Recherche Concertée” (ARC) (AUWB-2018-2023 ULB-No 7), Walloon Region grant (Win2Wal), FNRS Welbio grants (FNRS-WELBIO-CR-2017A-04 and FNRS-WELBIO-CR-2019A-04R), the FWO and FNRS under the Excellence of Science (EOS O.0020.22/RG3483) programme, the ULB Foundation, the Belgian Foundation against Cancer (FCC 2016-086 FAF-F/2016/872), and H2020-MSCA-ITN ROPES.

## AUTHOR CONTRIBUTIONS

Investigation, A.P., K.W., I.P., L.V.D.L., E. Calonne, B.H., C.H., P.P., L.L., Y.A., J.L., I.N.; Formal analysis, A.P., M.C., E. Collignon, M.B., F. M.; Visualization, A.P., M.C., E. Collignon; Writing, A.P., F.F.; Project Administration, S.L., P.J., C.Y., R.D., F.F., Funding acquisition, R.D., F.F.; Conceptualization and Supervision, F.F.

## METHODS

### LEAD CONTACT AND MATERIALS AVAILABILITY

Further information and requests for resources and reagents should be directed to and will be fulfilled by the lead contact François Fuks.

### EXPERIMENTAL MODEL AND STUDY PARTICIPANT DETAILS

#### Cell lines

HeLa (S3) cells were cultured in RPMI 1640 (Gibco) supplemented with 10% FBS (Gibco). HEK293GP cells were cultured in DMEM (Gibco) supplemented with 10% FBS (Gibco). All cells were grown at 37°C and humidified in a 5% CO2 atmosphere, are mycoplasma-free, and were authenticated. To generate HeLa cells devoid of mitochondrial DNA, cells were grown in media containing 50μg/mL uridine (Sigma) and 2.5μM ethidium bromide (Sigma), and passaged 8-10 times as previously described^2^.

#### Patient’s biopsies

Human cervical cancer samples were biobanked by Biobanque Hôpital Erasme-ULB (BE_BERA1), biobanque du laboratoire d’anatomie pathologique – CUB Hôpital Erasme; BBMRI-ERIC. The biobanque Hôpital Erasme-ULB, biobanque du service d’anatomie pathologique has been approved by the Ehics Committee of the Erasme-Université Libre de Bruxelles (Brussels, Belgium) in December 2018 (no. B2018/007) and has been notified to Federal Agency for Medicines and Health Products (no BB190015) in accordance with Belgian legal requirements.

## METHODS DETAILS

### CRISPR/Cas9-generated cell line

The METTL8 KO HeLa cell line was generated with the CRISPR/Cas9 nuclease system via homology-directed repair (HDR). Briefly, sgRNAs were designed to target the start codon (ATG) and about 500 bp downstream of ATG of METTL8 according to the guidelines listed on the CRISPOR website (http://crispor.tefor.net). The sgRNAs were cloned into the pX461 vector (Addgene, #48140). In parallel, donor pUC18 vectors (GenScript, #SD1162) were generated, containing gene sequences homologous to those flanking the sgRNA targeting sites and a bordering selectable marker (mCherry or puromycin resistance gene) and a mammalian transcriptional terminator (bGH, synthesized by GENEWIZ). Co-transfection of HeLa cells with the donor vectors and gRNA-containing plasmids at a ratio of 2:1 was performed with Lipofectamine™ 2000 according to the manufacturer’s instructions (Invitrogen). Twenty-four hours after transfection, selection was initiated with 2µg/ml puromycin and carried out for at least 10 days. Surviving mCherry-positive cells were sorted into 96-well plates (1 cell per well) by fluorescence-activated cell sorting (FACS) and grown for two to three weeks before clones were transferred into 24-well plates and METTL8 expression was measured by quantitative reverse transcription PCR (RT-qPCR) and western blotting. CRISPR-Cas9-targeted genomic regions in positive clones were PCR-amplified and sequenced. All relevant sgRNA sequences and primers are listed in Supplementary Table 1.

### Mitochondrial RNA isolation

Mitochondria were isolated using the Mitochondrial Isolation Kit (Abcam) according to the manufacturer’s recommendations. Supernatant was used to isolate the non-mitochondrial fraction. Total RNA extraction was performed with TRIzol and TRIzol LS (ThermoFischer) according to the manufacturer’s instructions. Genomic DNA was eliminated with the Ambion Dnase I (Rnase-free) kit. RNA was quantified and purity checked with the ND-1000 NanoDrop spectrophotometer (NanoDrop Technologies).

### RNA and DNA extraction

Total RNA was extracted with the Rneasy Kit (Qiagen). For mice tissues and xenografts, around 50mg of frozen samples were disrupted and homogenized using the TissueLyser LT (Qiagen) before RNA extraction using the Rneasy Lipid Tissue Mini Kit (Qiagen) according to the manufacturer’s instructions. Genomic DNA was eliminated by Dnase I treatment. DNA was extracted with the QIAamp DNA Mini Kit (Qiagen). DNA and RNA was quantified and purity checked with the ND-1000 NanoDrop spectrophotometer (NanoDrop Technologies).

### Poly(A) RNA enrichment

Poly(A) RNA enrichment was performed using the GenElute mRNA Purification Miniprep kit (Sigma) according to the manufacturer’s recommendations. Enrichment efficiency was assessed using the Bioanalyzer RNA 6000 Nano kit using the 2100 Bioanalyzer system (Agilent).

### LC-MS/MS for m^3^C detection and quantification

m^3^C detection by LC-MS/MS has been first described by Xu and colleagues^1^. 200 ng isolated RNA or 100 ng model RNA oligo was digested into nucleosides by 0.5U nuclease P1 (Sigma, N8630) in 20mL buffer containing 10mM ammonium acetate, pH 5.3 at 42°C for 6h, followed by the addition of 2.5mL 0.5M MES buffer, pH 6.5 and 0.5U alkaline phosphatase (Sigma, P4252). The mixture was incubated at 37°C for another 6h and diluted to 50μL. 5μL of the solution was injected into LC-MS/MS. The nucleosides were separated by ultra-performance liquid chromatography with a C18 column, and then detected by triplequadrupole mass spectrometer (AB SCIEX QTRAP 5500) in the positive ion multiple reactionmonitoring (MRM) mode.

### Reverse transcription and quantitative PCR

Isolated RNA was reverse-transcribed with the First Strand cDNA Synthesis kit (Roche) according to the manufacturer’s recommendations. Real-time PCR was performed with the LightCycler 480 SYBR Green I Master mix (Roche) on the LightCycler 480 real-time PCR system (Roche). Gene expression was normalized to human actin beta (ACTB) and succinate dehydrogenase complex flavoprotein subunit A (SDHA) or hypoxanthine phosphoribosyltransferase 1 (HPRT1). Primers are listed in Supplementary Table 1.

### Protein extraction and western blotting

Fresh cells, or mitochondria isolated using the Mitochondrial Isolation Kit (Abcam), were suspended in Lysis Buffer (Cell Signaling Technology) supplemented with Complete Protease and PhoStop phosphatase inhibitor cocktails (Roche). After centrifugation at 14,000g for 10 minutes at 4°C, proteins were quantified with the DC protein assay kit (Bio-Rad Laboratories). Cell extracts (25 to 50μg) were separated by SDS-PAGE and transferred to PVDF membranes (PerkinElmer Life Sciences). Membranes containing the transferred proteins were blocked with 5% nonfat dry milk with 0.1% tween-20 PBS solution for 1 hour, followed by overnight incubation with primary antibodies. For loading control, membranes were incubated with actin beta or vinculin antibodies. Immunocomplexes were detected with an ECL Plus system (Amersham Biosciences). Band densities were quantified with ImageJ software.

### Association between METTL8 expression and clinical features in cancer

TCGA data sets were downloaded directly from the Genomic Data Commons (GDC) website (https://gdc.cancer.gov/gdc-tcga-data-access-matrix-users)(data release 12.0). For GSEA, genes were pre-ranked using their Pearson correlation coefficients with METTL8 expression across all primary tumors (within each cancer type cohort). GSEA was then carried out in R (v4.3.0) with the fGSEA package (v.1.26.0). Most gene set collections were downloaded from the Molecular Signatures Database (v.7.5.1; http://www.gsea-msigdb.org/gsea/msigdb/index.jsp). For Fig. 2H, only ‘Hallmark’ gene sets significantly enriched (|NES| > 1.25 and adjusted p value < 0.05) in at least 5 TCGA cohorts are displayed in the heatmap. The custom mitochondrial gene signature was generated by randomly selecting 200 genes among those containing ’mitochondr’ in the ‘GOComponent’ column from the GRCh38.p13 genome annotation file (downloaded from GENCODE in July 2024; https://www.gencodegenes.org/human/release_38.html). Survival analyses were conducted using the Kaplan-Meier Plotter website (https://kmplot.com/) with the ’Auto select best cutoff’ setting, except for CESC, which was analyzed using TCGA data with the ‘survminer’ (v.0.4.9) and ‘survival’ (v.3.5-5) packages in R. The optimal cutoff for the expression of METTL8 was determined using the surv_cutpoint function from survminer, and survival differences were assessed using the log-rank test.

### Mitochondrial signal quantification in mouse tissues

To quantify the mitochondrial signal across different mouse tissues, we first retrieved tissue-specific gene expression data from the Human Protein Atlas (v.19, Ensembl v.92.38; https://v19.proteinatlas.org/about/download). The mitochondrial gene set was identified using GO terms related to mitochondria, as detailed above (see ‘Association between METTL8 expression and clinical features in cancer’) and a subset of 200 mitochondrial genes was selected for further analysis. We then calculated a mitochondrial signature score for each tissue by averaging the expression levels (nTPM) of the selected genes.

### METTL8 immunostaining in human tissues

Immunohistochemistry (IHC) images were retrieved from the Human Protein Atlas (https://v19.proteinatlas.org/), where METTL8 expression was measured using the HPA035421 antibody. IHC staining was performed on healthy tissues (available at https://v19.proteinatlas.org/ENSG00000123600-METTL8/tissue/primary+data) and various cancer types (including cervical cancer, accessible at https://v19.proteinatlas.org/ENSG00000123600-METTL8/pathology/cervical+cancer#ihc).

### m^3^C modified RNA immunoprecipitation (m^3^C-RIP)

The method for m^3^C-RIP was adapted from a protocol described previously^28^. Starting with 1mg total RNA, enrichment in polyadenylated RNA was done through one round of oligo-dT selection with the GenElute mRNA Miniprep kit (Sigma). The enriched RNA was then fragmented around 200- 300 nt by incubating 18μL RNA (at 0.9μg/μL) in 10μL of 10X fragmentation buffer (100mM Tris-HCl and 100mM ZnCl_2_) in thin-walled PCR tubes at 94°C for 18s in a pre-heated thermocycler block. Fragmentation was stopped by adding 2μL of 0.5M EDTA and the tubes were kept on ice. RNA was then precipitated with sodium acetate and resuspended in 300μL Rnase-free water. RNA fragment size was assessed on a 2100 Bioanalyzer (Agilent) with the RNA Nano kit (Agilent). 2-5μg of fragmented RNA was kept at -80°C to serve as input. The rest of RNA samples were denatured at 70°C for 5 min and then put on ice. Immunoprecipitation was then performed as follows: denatured fragmented poly(A) enriched RNA in 290μL, 10μL proteinase inhibitors (Roche), 5μL (200 units) Rnasin ribonuclease inhibitors (Promega), 5μL of 200nM Ribonucleoside Vanadyl Complex (RVC, Sigma), 100μL of 5X IP buffer (50mM Tris-HCl, 750mM NaCl, 0.5% (v:v) NP-40), 3.33μL anti-m^3^C antibody (Diagenode, C15410209), and 86.5μL Rnase-free water. The mix was incubated overnight at 4°C on a rotating wheel. The next day, 50μL of Dynabeads Protein G (Invitrogen) were washed twice with 1X IP buffer supplemented with antiproteases (Roche) and blocked with washing buffer supplemented with 0.5mg/mL BSA (Sigma) during 1h. Beads were washed again twice, added to the IP mix, and incubated for 2h at 4°C on a rotating wheel. The mix was washed 3 times with 1mL washing buffer supplemented with 10μL Rnasin and 10μL RVC. Elution was performed by TriPure (Roche) according to the manufacturer’s recommendations and resuspended in 9μL Rnase-free water. After RNA recovery, cDNA libraries were prepared with the SMARTer stranded total RNA-Seq kit v2 – pico input mammalian (Takara) for both input and IP samples. Sequencing was performed with the Illumina NextSeq500 (Illumina). For m^3^C-RIP-qPCR, 1mg of total RNA was immunoprecipitated without poly(A) enrichment, according to the above method. Quantitative PCR was then performed using *tRNA-Ser*, *tRNA-Thr*, *tRNA-Arg* and *tRNA-Asn* primers (primers available in Supplementary Table 1). All experiments were performed at least in biological duplicates.

### Reads pre-processing

Sequencing data were pre-processed in the following steps. First, the raw sequencing data were analyzed with FastQC (v0.11.5)^29^. Low-complexity reads were removed with the AfterQC tool (v0.9.6)^30^ with default parameters. To get rid of reads originating from rRNA or tRNA, reads were mapped with Bowtie2 (v2.3.4.1)^31,32^ to Homo Sapiens tRNA and rRNA sequences were downloaded from https://www.ncbi.nlm.nih.gov/nuccore using “Homo sapiens”[Organism] AND (biomol_rrna [PROP] OR biomol_trna [PROP] as search parameters. Unmapped reads were then further processed with Trimmomatic (v0.33)^32^ using default parameters to remove adapter sequences. The resulting fastq data were again analyzed with FastQC to ensure that no further processing was needed.

### m^3^C-RIP-seq: mitochondrial analysis

The pre-processed reads were mapped against the human reference genome (GRCh37/hg19) with the STAR algorithm (v2.6.0c)^33^ using the reference transcriptome based on Ensembl (v85)^34^ and LNCipedia (v5.2)^35^. For each condition, replicates were pooled to form a metasample. Mitochondrial m^3^C peak regions were identified by applying the m^6^Aviewer peak-calling tool (v1.6.1)^36^ on mitochondrial reads. Sites showing significant enrichment over input in metasample and showing a differential enrichment between WT and METTL8 KO were considered. Peaks were resized to 100 bp on both sides of the identified summit. To obtain visual representations of local enrichment profiles, normalized (HPB) bedgraph files were generated using bamTobw (https://github.com/YangLab/bamTobw)^37^ and uploaded into the IGV tool (v2.9.4)^38^.

### Mitochondrial content determination

Mitochondria DNA copy number was estimated as previously described^39^. Briefly, quantitative real-time polymerase chain reaction (qPCR) was performed with the LightCycler 480 SYBR Green I Master mix (Roche) on the LightCycler 480 real-time PCR system (Roche) using the mitochondrial cytochrome b (*CYTB*) and the single-copy nuclear β-2 microglobulin (*B2M*) gene (primers available in Supplementary Table 1). Quantification of mt-DNA was accomplished by calculating the ratio of mt- DNA-encoded *CYTB* gene to the nDNA-encoded *B2M* gene.

The fluorescent labeling of mitochondria was performed with the MitoTracker Green FM (Thermo Fisher), according to the manufacturer’s recommendations. Briefly, 24h after cells seeding (200,000 cells per well) in a 24-wells plate, cells were washed 2 times with PBS. Cells were then incubated in a medium without serum containing the MitoTracker Green FM (1:1000) and the Incucyte Nuclight rapid red dye for live-cell nuclear labeling (Essen Bioscience) (1:1000) for 30 min at 37°C. Imaging and quantification was performed using an IncuCyte S3 machine (Essen Bioscience) and the accompanying commercial software.

### Complex I activity measurement

Complex I activity was measured in cells with the complex I enzyme activity microplate assay kit (Abcam) according to the manufacturer’s recommendations. Relative activity is expressed as the change in absorbance per minute (mOD min^-1^).

### Metabolomics

For metabolomic analysis the extraction solution was composed of 50% methanol, 30% acetonitrile (I) and 20% water. The volume of extraction solution was adjusted to cell number (1ml per 10^6^ cells). After addition of extraction solution, samples were vortexed for 5min at 4 °C and centrifuged at 16,000g for 15min at 4°C. The supernatants were collected and stored at −80° C until analysis. LC/MS analyses were conducted on a QExactive Plus Orbitrap mass spectrometer equipped with an Ion Max source and a HESI II probe coupled to a Dionex UltiMate 3000 uHPLC system (ThermoFisher). External mass calibration was performed using a standard calibration mixture every seven days, as recommended by the manufacturer. The 5µl samples were injected onto a ZIC-pHILIC column (15 mm × 2.1mm; i.d. 5µm) with a guard column (20mm × 2.1mm; i.d. 5µm) (Millipore) for LC separation. Buffer A was 20mM ammonium carbonate, 0.1% ammonium hydroxide (pH 9.2), and buffer B was ACN. The chromatographic gradient was run at a flow rate of 0.200µl min−1 as follows: 0–20min, linear gradient from 80% to 20% of buffer B; 20–20.5min, linear gradient from 20% to 80% of buffer B; 20.5–28min, 80% buffer B. The mass spectrometer was operated in full scan, polarity switching mode with the spray voltage set to 2.5kV and the heated capillary held at 320°C. The sheath gas flow was set to 20 units, the auxiliary gas flow to 5 units and the sweep gas flow to 0 units. The metabolites were detected across a mass range of 75–1,000 m/z at a resolution of 35,000 (at 200 m/z) with the automatic gain control target at 106 and the maximum injection time at 250ms. Lock masses were used to ensure mass accuracy below 5ppm. Data were acquired with Thermo Xcalibur software (ThermoFisher). The peak areas of metabolites were determined using Thermo TraceFinder software (ThermoFisher), identified by the exact mass of each singly charged ion and by the known retention time on the HPLC column. All data were analyzed with the MetaboAnalyst interface and are available in Table S2.

### Proliferation and migration assays

WT and METTL8 knockout cells (1,000 cells per well) were seeded in a 96-well plate (Corning) and transferred to the Incucyte S3 live-cell analysis instrument (Essen Bioscience). Growth profile was monitored by 10X objective every 2h using the Incucyte software for 5 days. For migration assay, WT and METTL8 knockout cells (45,000 cells per well) were seeded in a 96-well ImageLock tissue culture plate (Essen Bioscience). The next day, wounds were created in all wells using the Woundmaker (Essen Bioscience) and plates were transferred to the Incucyte S3 live-cell analysis instrument (Essen Bioscience). Migration profile was monitored by 4X objective every hour using the Incucyte software for 24h. Standard mode per well was used to collect images in phase-contrast mode and averaged to provide a representative statistical measure of the well confluency. All experiments included technical replicates and were performed at least three independent times. Results are shown as the mean of the triplicate.

### Mouse xenografts

HeLa WT and METTL8 knockout cells were injected (5 x 10^6^ cells per flank in 100μL of PBS with 20% BD Matrigel matrix) subcutaneously in female athymic nude mice (Charles River) of 4 weeks of age (5 mice per condition). Tumor growth was monitored every week for tumor appearance and progression.

For luciferase experiments, HeLa WT and METTL8 knockout cells were engineered to stably express firefly luciferase. Briefly, the retroviral transduction of HeLa WT and METTL8 KO cells with the MSCV luciferase PGK-hygro (plasmid was a gift from Scott Lowe, Addgene plasmid #18782) was performed by transfecting 293GP cells with the MSCV luciferase PGK-hygro plasmid and pVSV-g plasmid and using the subsequent viral supernatant to infect the HeLa cells.

Luciferase expressing HeLa WT and METTL8 KO cells were injected (1.5 x 10^6^ cells per flank in 100μL of PBS with 20% BD Matrigel matrix) subcutaneously in female athymic nude mice (Charles River) of 4 weeks of age (5 mice per condition). Following subcutaneous injection of 150mg/kg of D-luciferin (Promega), tumor growth was monitored every 3-4 days by bioluminescence imaging (BLI) in a PhotonImager Optima (Biospace Lab), which dynamically counted the emitted photons for 25min, while animals are under anesthesia (4% and 2.2% isoflurane for initiation and maintenance, respectively). Image analysis was performed with M3Vision software (Biospace Lab). At the end of experiments, xenografts were dissected and biopsies were snap-frozen in liquid nitrogen for further analysis. Animal experiments were performed under approval of local Animal Ethics Evaluation Committee (CEBEA protocol No: CEBEA- CMMI-2021-02).

### RNA-seq and data analyses

Following RNA isolation, previously described, cDNA libraries were constructed with 500ng total RNA using the Illumina TruSeq stranded total RNA ribo-zero H/M/R gold kit (Illumina) according to the manufacturer’s recommendations (5 xenografts per condition). Libraries qualities were assessed and quantified with a 2100 Bioanalyzer (Agilent) using the high sensitivity DNA kit (Agilent), and Qubit 2.0 (ThermoFisher). Sequencing was performed with the Illumina NextSeq500 (Illumina). Sequencing reads were processed as described under ‘Reads pre-processing’. To exclude mouse transcript contamination, the pre-processed reads were mapped against the mouse reference genome (mm9) with the STAR algorithm (v2.6.0c) using Ensembl transcriptome (v67). The unmapped reads were then mapped against the human reference genome (GRCh37/hg19) with the STAR algorithm (v2.6.0c) using the reference transcriptome based on Ensembl (v85)^34^ and LNCipedia (v5.2)^35^. Gene expression was computed with the HTseq tool (v0.9.1)^36^. Raw gene expression counts were subjected to DESeq2 (v1.22.2)^40^ for normalization and analysis of differential expression between METTL8 KO and WT xenografts. Genes were reported as differentially expressed if a log fold change of at least 1 was observed between the two conditions and the p-value adjusted was below 0.05. Normalized counts were log-transformed and mean-centered for PCA and heatmap representations, and hierarchical clustering was performed using the ’ward.D’ method. Similar to TCGA analyses, GSEA was carried out in R (v4.3.0) with the fGSEA package (v.1.26.0), with genes being pre-ranked by log_2_ fold change from the differential expression analysis (METTL8 KO/WT). Gene set collections were downloaded from the Molecular Signatures Database (v.7.5.1; http://www.gsea-msigdb.org/gsea/msigdb/index.jsp).

## QUANTIFICATION AND STATISTICAL ANALYSIS

Statistical analyses were performed using either the computing environment R or GraphPad Prism 9 software, and results are expressed as mean±SEM or mean±SD. All statistical details of experiments can be found in the figure legends. For 2-group comparison, 2-tailed unpaired *t* test or Mann-Whitney test was performed. For multiple-group comparison, 1-way ANOVA or Kruskal-Wallis tests were performed with the multiple-comparison post hoc correction as indicated. Equality-of-variance test between groups and Shapiro-Wilk normality test were performed, and statistical tests were chosen accordingly. Graphs show exact *p*-values and *p* less than 0.05 was considered significant. When box plots are used to visualize data distribution, whiskers are down to the minimum and up to the maximum value, and each individual value are plotted as a point. Otherwise indicated, all experiments included technical replicates and were performed at least three independent times.

## SUPPLEMENTAL TABLES

**Table S1. List of primers**

**Table S2. Metabolomics**

## SUPPLEMENTAL FIGURES

**Supplemental Figure 1 (related to Figure 1).**
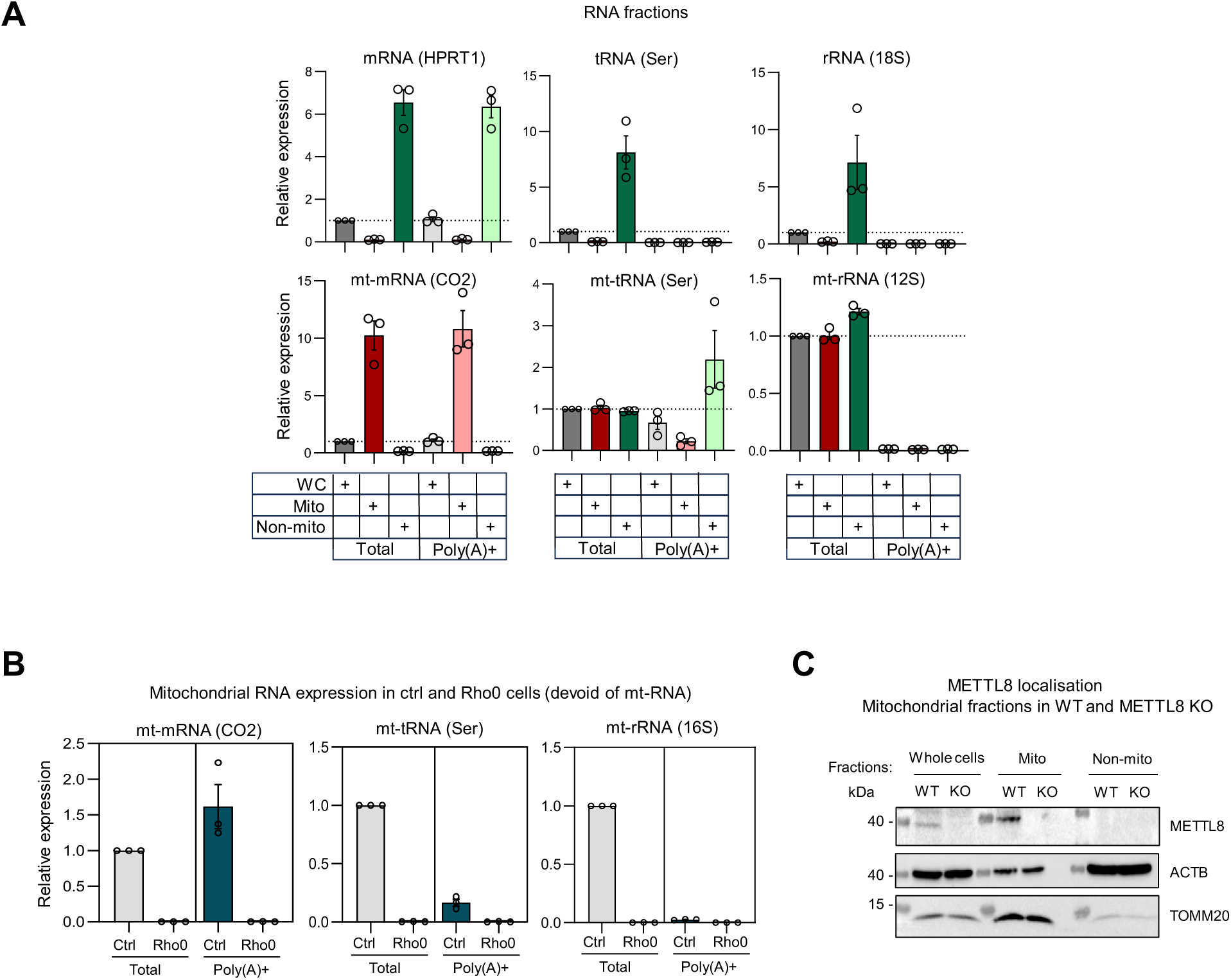
METTL8-dependent m^3^C on mt-mRNA. (A) Relative quantification of mRNA (HPRT1), tRNA (Ser), rRNA (18S), mt-mRNA (CO2), mt-tRNA (Ser) and mt-rRNA (12S) transcripts from the total RNA and poly(A) RNA from whole-cell (WC), mitochondrial and non-mitochondrial fractions. (n=3, mean± SEM) (B) Relative quantification of mt-mRNA (CO2), mt-tRNA (Ser), and mt-rRNA (16S) transcripts from control and Rho0 cells, in total RNA and poly(A) RNA (n=3, mean± SEM). (C) Representative western blot image of METTL8 in whole-cell (WC), mitochondrial and non-mitochondrial fractions, in WT and METTL8 KO HeLa cells (*n*=3).

**Supplemental Figure 2 (related to Figure 2).**
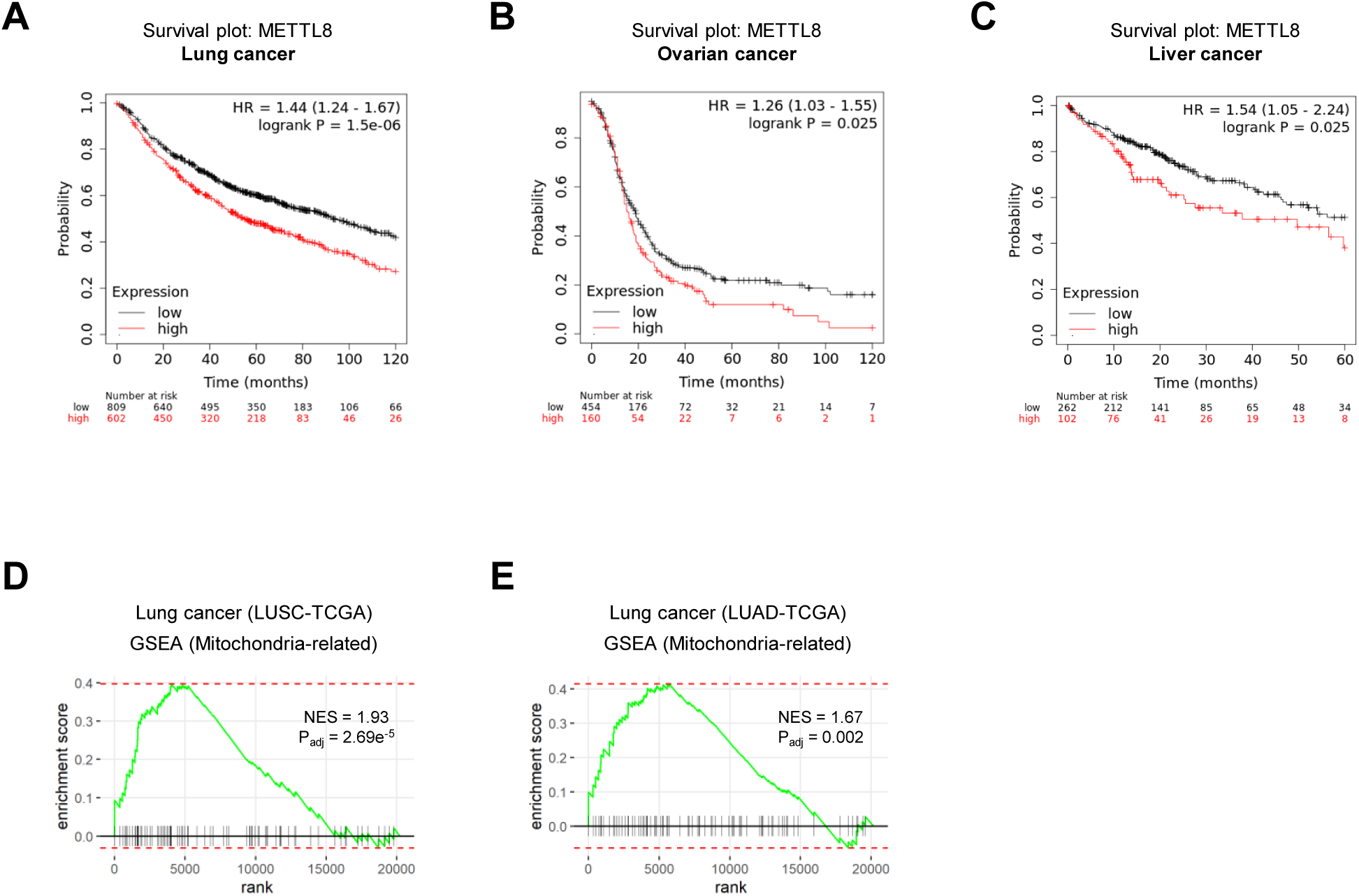
METTL8 expression and signatures in cancers. (A-C) Kaplan-Meier curves showing the overall survival of patients categorized as METTL8 high or low in (A) lung cancer, (B) ovarian cancer, (C) liver cancer. (Hazard Ratio (HR) with 95% confidence interval and logrank test) (D-E) GSEA in TCGA lung cancer (D, LUSC; E, LUAD) using a signature of mitochondria-related genes. All genes were pre-ranked based on correlation with METTL8 expression.

**Supplemental Figure 3 (related to Figure 3).**
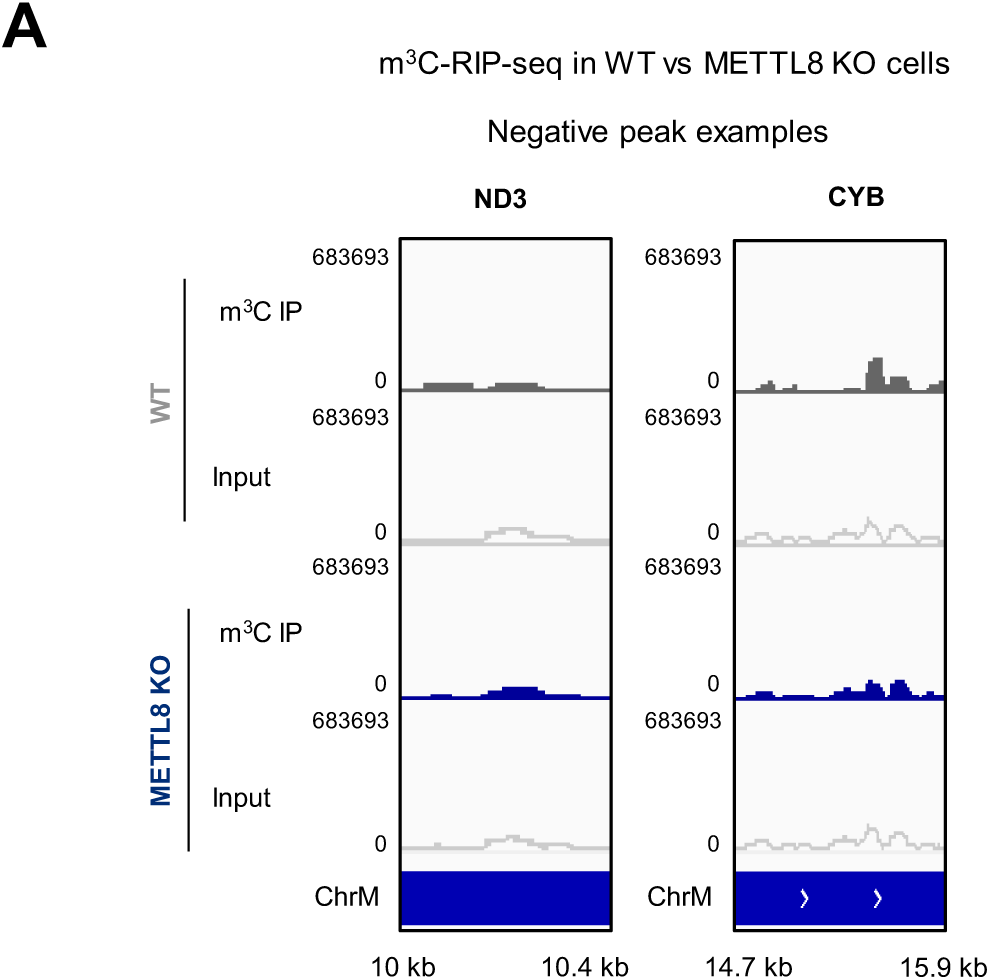
mt-mRNA m^3^C mapping. (A) Example tracks of negative peaks of m^3^C by RIP-seq (ND3 and CYB) in WT vs METTL8 KO HeLa cells.

**Supplemental Figure 4 (related to Figure 4).**
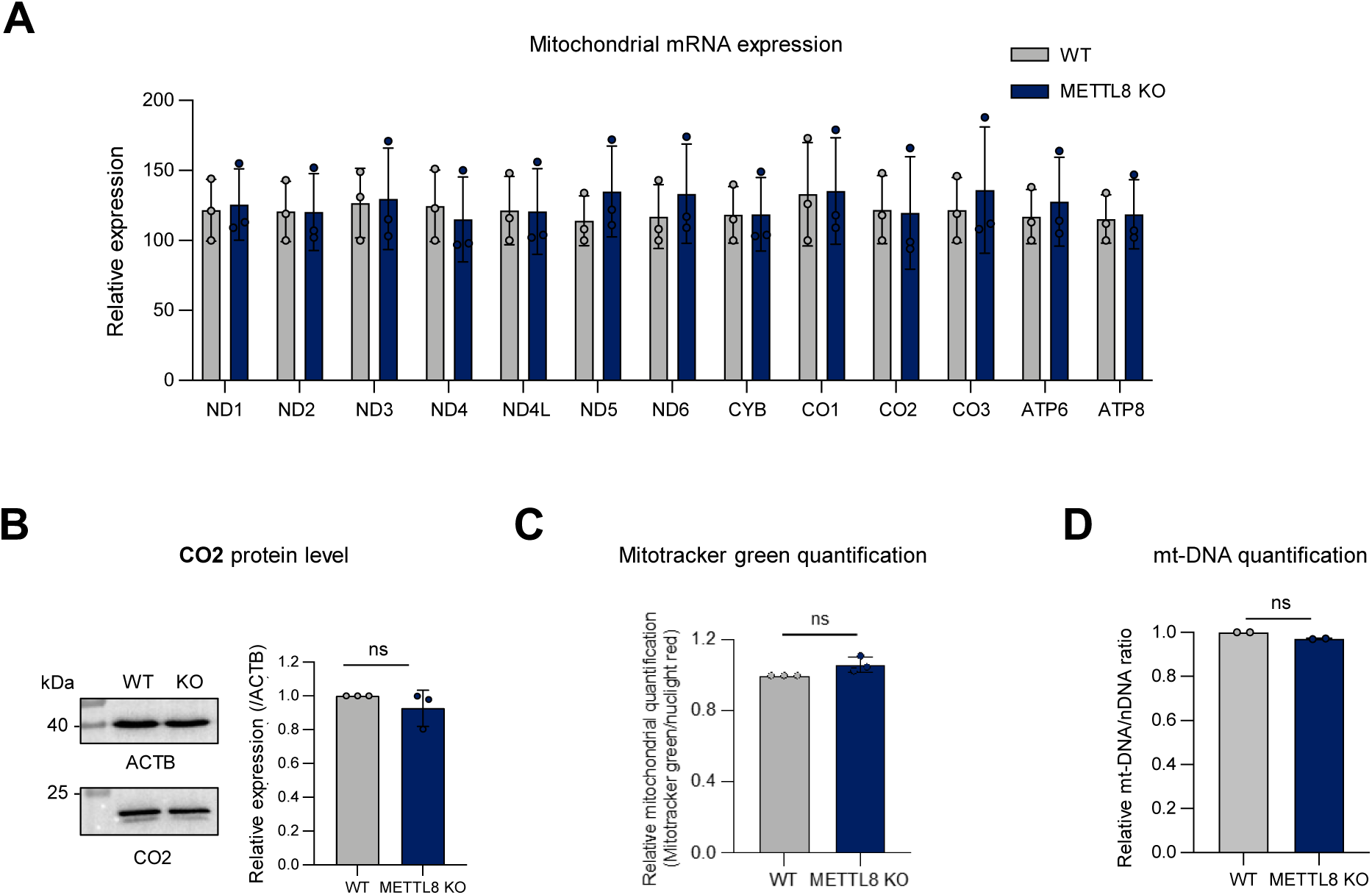
Roles of METTL8-dependent m^3^C. (A) Relative quantification of all mt-mRNA in WT vs METTL8 KO cells (*n*=3, mean±SD). (B) Representative western blot image of CO2 in WT and METTL8 KO cells with quantification of protein levels (normalized to ACTB, *n*=3, two-tailed *t* test, mean±SEM). (C) Relative mitochondrial quantification. Ratio between Mitotracker Green signal (mitochondrial staining) and Nuclight Red signal (nuclear staining) in WT and METTL8 KO cells (two-tailed *t* test, ns: not significant). (D) Relative quantification of mitochondrial DNA in WT and METTL8 KO cells, expressed as the CYTB/B2M ratio (*n*=2, two-tailed *t* test, ns: not significant).

**Supplemental Figure 5 (related to Figure 5).**
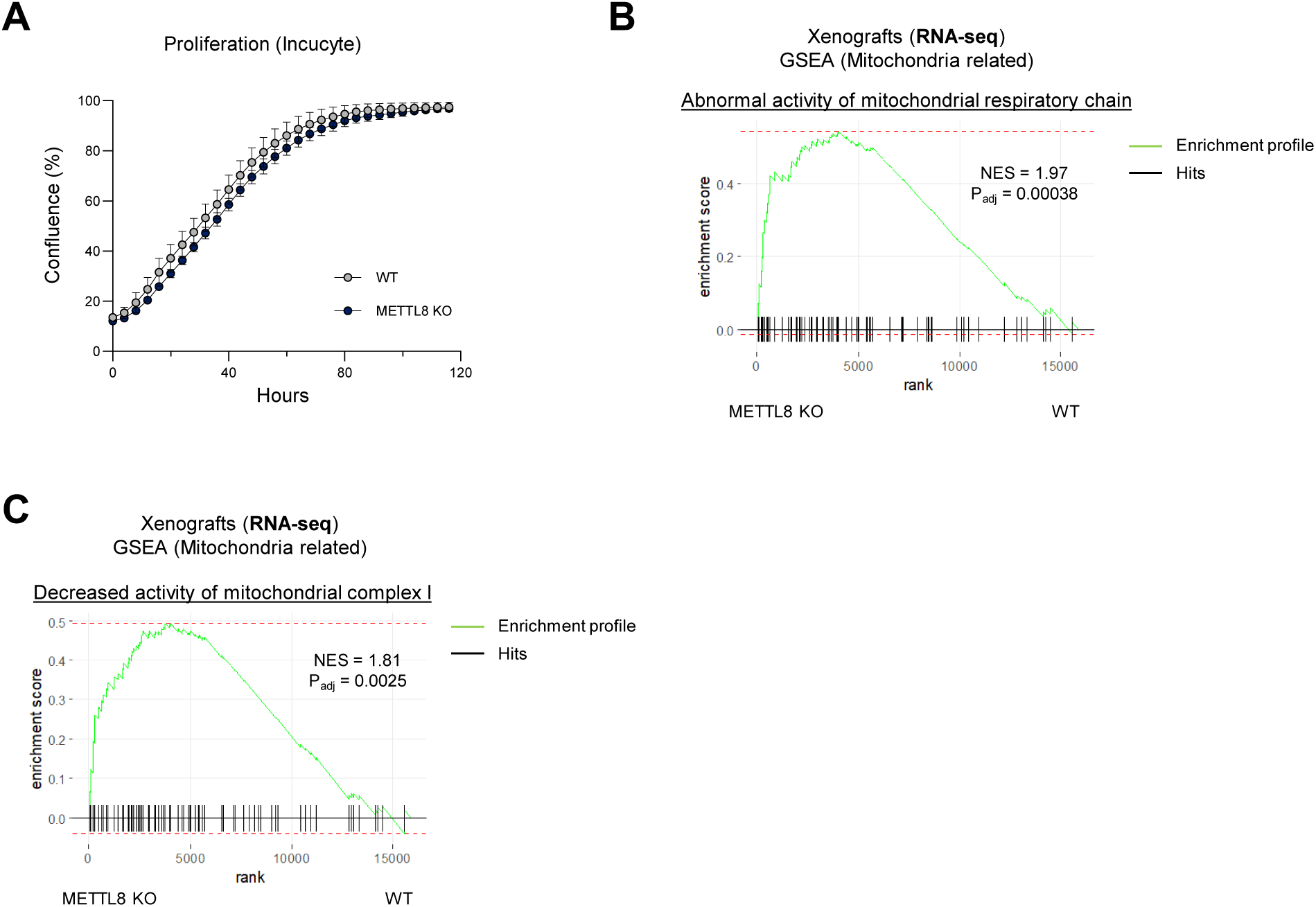
METTL8 and cervical cancer. (A) Real-time proliferation of WT and METTL8 KO cells (*n*=3, mean±SEM). (B-C) GSEA in xenograft RNA-seq data, using the ‘Abnormal activity of mitochondrial respiratory chain’ and ‘Decreased activity of mitochondrial complex I’ gene sets. Genes were pre-ranked based on differential expression (fold change) upon METTL8 depletion.

