## Supplementary material for "m^3^C is a mitochondrial mRNA modification which promotes tumor progression": Table S1

**Supplementary table 1. Primers and oligos sequences**

| Primers sequences for METTL8 CRISPR/Cas9-generated HeLa cells | | |
|---|---|---|
| ***METTL8* gRNA A** | Forward | 5'-CACCGAAATTCCATTTCTTGTCTA-3' |
| | Reverse | 5'-AAACTAGACAAGAAATGGAATTTC-3' |
| ***METTL8* gRNA B** | Forward | 5'-CACCTAACCACTTTGGTATCTGTG-3 |
| | Reverse | 5'-AAACCACAGATACCAAAGTGGTTA-3' |
| ***METTL8* gRNA C** | Forward | 5'-CACCAGCTGATAATTACCCCTACA-3' |
| | Reverse | 5'-AAACTGTAGGGGTAATTATCAGCTC-3' |
| ***METTL8* gRNA D** | Forward | 5'-CACCATCTATCTTTCCAAATAGGG-3' |
| | Reverse | 5'-AAACCCCTATTTGGAAAGATAGAT-3' |
| **Left homologous arm** | Forward | 5'-GATTACGAATTCCTTCCAGAAATGGAGATGGATTTGG-3' |
| | Reverse mCherry | 5'-GCCCTTGCTCACCATCCTAAGCAGTACCTAAAAAG-3' |
| | Reverse puromycin resistance | 5'-CTTGTACTCGGTCATCCTAAGCAGTACCTAAAAAG-3' |
| **Right homologous arm** | Forward | 5'-GCATGCTGGGGAGCAGTAAATTGTGAGGGCTCAAT-3' |
| | Reverse | 5'-AGTGCCAAGCTTGAATTCCTTGCAAAATACTACTGCT-3' |
| **mCherry + terminator gene** | Forward | 5'-GTACTGCTTAGGATGGTGAGCAAGGGCGAGGAGGATAAC-3' |
| | Reverse | 5'-ACAATTTACTGCTCCCCAGCATGCCTGCTATTCTCTT-3' |
| **Puromycine resistance + terminator gene** | Forward | 5'-GTACTGCTTAGGATGACCGAGTACAAGCCCACGGTGC-3' |
| | Reverse | 5'-ACAATTTACTGCTCCCCAGCATGCCTGCTATTCTCTT-3' |
| **Plasmid pUC18** | Forward | 5'-TGCAAGGAATTCAAGCTTGGCACTGGCCGTCGTTTTAC-3' |
| | Reverse | 5'-CATTTCTGGAAGGAATTCGTAATCATGGTCATAGCT-3' |

| Primers sequences for Mettl8 CRISPR/Cas9-generated mice | | |
|---|---|---|
| **Mettl8 crRNA-1** | | 5'-GACAATCAACGTCAGACTGT-3' |
| **Mettl8 crRNA-2** | | 5'-GACTAACACCATATGAACCG-3' |
| **Mettl8 genotyping** | Forward | 5'-GCACTGCCTCATCACAGAAAC-3' |
| | Reverse | 5'-TTGGTTTACACATTGCCCACC-3' |

| Primers sequences for RT-qPCR | | |
|---|---|---|
| ***12S*** | Forward | 5'-ACTGCTCGCCAGAACACTAC-3' |
| | Reverse | 5'-GGTGAGGTTGATCGGGGTTT-3' |
| ***16S*** | Forward | 5'-ATGAATGGCTCCACGAGGG-3' |
| | Reverse | 5'-CTTGCTGTGTTATGCCCGC-3' |
| ***18S*** | Forward | 5'-CGCTACTACCGATTGGATGG-3' |
| | Reverse | 5'-ACCTTGTTACGACTTTTACTT-3' |
| ***28S*** | Forward | 5'-TGCCCAGTGCTCTGAATGTC-3' |
| | Reverse | 5'-ACTCCCGCCGTTTACCCG-3' |
| ***ACTB*** | Forward | 5'-TAATGTCACGCACGATTTCC-3' |
| | Reverse | 5'-CGGGACCTGACTGACTACCT-3' |
| ***B2M*** | Forward | 5'-CCAGCAGAGAATGGAAAGTCAA-3' |
| | Reverse | 5'-TCTCTCTCCATTCTTCAGTAAGTCAACT-3' |
| ***CYTB*** | Forward | 5'-GCCTGCCTGATCCTCCAAAT-3' |
| | Reverse | 5'-AAGGTAGCGGATGATTCAGCC-3' |
| ***HPRT1*** | Forward | 5'-TGACACTGGCAAAACAATGCA-3' |
| | Reverse | 5'-GGTCCTTTTCACCAGCAAGCT-3' |
| ***METTL8*** | Forward | 5'-ACAAAAGGGGAAGTCCACAGT-3' |
| | Reverse | 5'-TTGTAAGCGGCGATCAACCA-3' |
| ***MT-ATP6*** | Forward | 5'-TGCCACAACTAACCTCCTCG-3' |
| | Reverse | 5'-GGATGGCCATGGCTAGGTTT-3' |
| ***MT-ATP8*** | Forward | 5'-ACTACCACCTACCTCCCTCAC-3' |
| | Reverse | 5'-GGATTGTGGGGGCAATGAATG-3' |
| ***MT-CO1*** | Forward | 5'-CCTATCATCTGTAGGCTCATTC-3' |
| | Reverse | 5'-GGAGGGTTCTTCTACTATTAGGAC-3' |
| ***MT-CO2*** | Forward | 5'-CTGCGACTCCTTGAGCTTGA-3' |
| | Reverse | 5'-TCGTGTAGCGGTGAAAGTGG-3' |
| ***MT-CO3*** | Forward | 5'-CGCCTGATACTGGCATTTTG-3' |
| | Reverse | 5'-GACCCTCATCAATAGATGGAGAC-3' |
| ***MT-CYB*** | Forward | 5'-ACCCCCTAGGAATCACCTCC-3' |
| | Reverse | 5'-GCCTAGGAGGTCTGGTGAGA-3' |
| ***MT-ND1*** | Forward | 5'-CGAACAGCATACCCCCGATT-3' |
| | Reverse | 5'-TGCTAGGGTGAGTGGTAGGA-3' |
| ***MT-ND2*** | Forward | 5'-ACCATCTTTGCAGGCACACT-3' |
| | Reverse | 5'-GCTTCTGTGGAACGAGGGTT-3' |
| ***MT-ND3*** | Forward | 5'-CCGCGTCCCTTTCTCCATAA-3' |
| | Reverse | 5'-GGCCAGACTTAGGGCTAGGA-3' |
| ***MT-ND4*** | Forward | 5'-CCCTCGTAGTAACAGCCATTCTC-3' |
| | Reverse | 5'-GACTGTGAGTGCGTTCGTAGT-3' |
| ***MT-ND4L*** | Forward | 5'-TCGCTCACACCTCATATCCTC-3' |
| | Reverse | 5'-AGGCGGCAAAGACTAGTATGG-3' |
| ***MT-ND5*** | Forward | 5'-CACATCTGTACCCACGCCTT-3' |
| | Reverse | 5'-AATGCTAGGCTGCCAATGGT-3' |
| ***MT-ND6*** | Forward | 5'-GGGAGGATCCTATTGGTGCG-3' |
| | Reverse | 5'-CCTATTCCCCCGAGCAATCT-3' |
| ***MT-tRNA Ser*** | Forward | 5'-GAAAAAGTCATGGAGGC-3' |
| | Reverse | 5'-ACCCCCCAAAGCTGGTT-3' |
| ***MT-tRNA Thr*** | Forward | 5'-AAACTAATACACCAGTCTTGTAAACC-3' |
| | Reverse | 5'-CCTTGGAAAAAGGTTTTCATCT-3' |
| ***SDHA*** | Forward | 5'-TGGTGCTGGTTGTCTCATTA-3' |
| | Reverse | 5'-ACCTTTCGCCTTGACTGTT-3' |
| ***tRNA Arg*** | Forward | 5'-CGGTGGCCTAATGGATAA-3' |
| | Reverse | 5'-GGGACTCGAACCCACAATCC-3' |
| ***tRNA Asn*** | Forward | 5'-GTCTCTGTGGCGCAATCGG-3' |
| | Reverse | 5'-CGAACCGCCAACCTTTTAGTTA-3' |
| ***tRNA Ser*** | Forward | 5'-CCGAGTGGTTAAGGCGATGG-3' |
| | Reverse | 5'-GAAACCCCAATGGATTTCAA-3' |
| ***tRNA Thr*** | Forward | 5'-CGTGGCTTAGTTGGTTAAAG-3' |
| | Reverse | 5'-GGATCTCCTGTTTACTAGACAG-3' |
