## Supplementary material for "m^3^C is a mitochondrial mRNA modification which promotes tumor progression": Table S2

**Table S2. Metabolomics**

| | HeLa | | | HeLa | | |
|---|---|---|---|---|---|---|
| | WT #1 | WT #2 | WT #3 | METTL8 KO #1 | METTL8 KO #2 | METTL8 KO #3 |
| 3-hydroxybutyric acid | 0,11377426 | 0,8744892 | 1,04810997 | -0,035211361 | -0,252174433 | -1,748987638 |
| 5-oxo-L-Proline | 0,08029493 | 1,66054769 | -0,85908188 | 0,151224082 | -1,202682941 | 0,169698121 |
| 6-Phosphogluconic acid | 0,98682953 | 1,01618215 | -1,01552828 | -0,863078258 | -0,839891845 | 0,715486707 |
| Acetyl CoA | 1,0047442 | 0,89464184 | 0,5865488 | -1,13276576 | -1,245519522 | -0,10764956 |
| acetyl-aspartate (N) | 0,75736809 | -0,06789919 | 1,5012093 | -0,539452452 | -0,318724478 | -1,332501279 |
| acetyl-carnitine | 0,70760762 | 0,79727984 | 1,18411872 | -0,958113298 | -1,01631558 | -0,714577301 |
| acetyl-glutamine | 1,44015173 | 0,50978723 | -0,67720097 | -0,303506431 | 0,422483382 | -1,39171494 |
| acetyl-lysine | -0,67610781 | -0,4430154 | -1,33359626 | 0,225753554 | 0,976582743 | 1,250383174 |
| Adenine | 0,35018841 | -0,3069278 | 1,37038811 | -1,618028018 | 0,467086707 | -0,262707414 |
| Adenosine | -1,20489252 | -0,49377341 | 1,37133744 | 0,033104176 | -0,683392386 | 0,977616704 |
| ADP | 0,79492589 | 1,06688301 | -0,2001499 | -1,531277669 | -0,711906423 | 0,581525099 |
| aKG | -0,25966922 | 0,70537003 | 1,4020338 | 0,187273954 | -0,591109642 | -1,44389892 |
| Aminoadipate | 1,46244846 | 0,61920195 | 0,49811294 | -0,838893667 | -0,797425436 | -0,943444245 |
| AMP | 0,63634464 | 0,91350849 | 1,13882979 | -1,032042436 | -0,942980674 | -0,713659803 |
| Arachidonic acid | 0,89647613 | 1,45608042 | -0,10195454 | -1,030775484 | -0,249858296 | -0,969968227 |
| Arginine | -0,82140491 | 0,45645959 | -0,89870543 | 1,423009072 | 0,717696164 | -0,877054482 |
| Argininosuccinate | 0,11652355 | -0,58530701 | -1,43003599 | 0,481445588 | 1,53408709 | -0,116713235 |
| Asparagine | -0,4850312 | -1,19600846 | -0,85937435 | 0,745868987 | 1,360558484 | 0,43398653 |
| Aspartate | 0,55157833 | -0,15062639 | 0,678227 | -0,406743173 | 1,046091097 | -1,718526863 |
| ATP | 0,82004067 | 0,26333705 | -0,67311683 | -1,321668947 | -0,450651172 | 1,362059226 |
| beta-Alanine | -1,05554791 | -1,26815838 | 0,65332617 | 1,234690215 | -0,12229109 | 0,557980996 |
| betaine | -1,67373854 | 0,47063106 | -0,11327277 | 1,261515879 | 0,458331033 | -0,40346666 |
| Butyric acid | 0,49611823 | 1,60801511 | -0,13511526 | -0,329797987 | -0,22901288 | -1,410207207 |
| Butyryl-carnitine | 0,99901106 | -0,16643068 | 1,40050189 | -0,785390511 | -1,141055792 | -0,306635968 |
| carnitine | -0,59841005 | -1,2225884 | -0,70144608 | 0,866676318 | 0,311529373 | 1,344238831 |
| Carnosine | 0,80467747 | 0,87573619 | -1,54618729 | 0,748201579 | -0,090685795 | -0,791742153 |
| CDP | 1,12518805 | 0,44631395 | -0,9366449 | -0,979481583 | -0,732863111 | 1,077487598 |
| cis-aconitate | 0,16701947 | 0,91400494 | 1,05085819 | 0,218137344 | -1,509312025 | -0,84070793 |
| Citrate | 0,18251093 | 1,27322886 | -0,43302363 | -0,551331857 | -1,406802775 | 0,935418459 |
| Citrulline | -0,51105922 | -0,26615278 | -0,37376652 | 2,032297828 | -0,371077208 | -0,510242106 |
| CMP | -0,9498751 | 1,18476426 | 1,28945504 | -0,162199282 | -0,87773038 | -0,484414533 |
| Creatine | 0,59786034 | 0,27127606 | 1,54503415 | -0,586018232 | -0,624806436 | -1,203345889 |
| Creatinine | -1,10140675 | 0,97989899 | -1,29546371 | 0,903613917 | -0,059698212 | 0,573055775 |
| CTP | 0,21941969 | 0,34618705 | -0,86576223 | -1,109501591 | -0,25899326 | 1,668650351 |
| Cystathionine | -0,07269929 | 0,1369079 | -1,44341868 | 0,54362144 | 1,477955881 | -0,642367256 |
| Cysteinesulfinic acid | 0,03272643 | -0,93499924 | 0,13290514 | 1,157327656 | 0,966273824 | -1,354233813 |
| Cystine | 1,60785839 | -0,88005725 | 0,49693377 | 0,289364632 | -1,042852883 | -0,471246661 |
| Decanoic acid | 0,09054918 | -0,25355834 | 1,74661636 | 0,060072048 | -0,310766761 | -1,332912481 |
| Decanoyl-Carnitine | -0,22183653 | 0,28953395 | 1,74762068 | -1,279317911 | -0,395432794 | -0,140567395 |
| dihydroxyacetone phosphate | 0,70474296 | 0,62633848 | 1,31705089 | -0,974647078 | -0,722285168 | -0,951200093 |
| Docosahexaenoic acid | 0,65035217 | -1,13430142 | 1,30381622 | 0,339289556 | -1,213426156 | 0,054269631 |
| Dodecanoyl-carnitine | 0,15921536 | 0,63934306 | 1,50853782 | -1,086969967 | -0,183245366 | -1,036880902 |
| FAD | 1,21634359 | 0,56329807 | 0,68296546 | -1,504802099 | -0,561477134 | -0,39632789 |
| Fructose | -0,07816721 | -0,25946025 | -1,77946825 | 0,547106515 | 0,447130382 | 1,122858807 |
| Fumarate | 1,37044389 | 0,18024012 | 0,46561044 | -1,666384028 | -0,306388799 | -0,043521622 |
| G6P | 0,99485923 | 1,16018323 | -0,45243991 | -1,430903829 | -0,568938408 | 0,297239686 |
| GDP | 0,91906078 | 1,0631359 | -1,02485074 | -1,028323804 | -0,640817775 | 0,711795647 |
| GLN | -0,87109211 | -0,85623886 | -0,98643123 | 0,997847937 | 1,041158646 | 0,674755617 |
| GLU | -0,82237437 | -1,05385297 | -0,85344803 | 0,931426822 | 0,886555753 | 0,911692793 |
| Glucose | -0,1597634 | 0,37579433 | -1,58149327 | 1,443358353 | 0,31163442 | -0,389530433 |
| Glyceraldehyde 3-phosphate | 0,32478231 | -0,59785508 | -1,34335893 | 1,415973279 | -0,4944978 | 0,694956215 |
| Glycerol 3-phosphate | 0,75680555 | 1,23646251 | 0,46878781 | -1,363694968 | -0,877647715 | -0,220713197 |
| Glycine | -0,94608801 | -1,14485579 | 1,55226836 | -0,232253565 | 0,25589179 | 0,515037205 |
| GMP | -0,83041192 | 1,16168901 | 0,1493361 | -0,85114294 | -0,850472297 | 1,221002042 |

| | | | | | | |
|---|---|---|---|---|---|---|
| GSH | 0,80969118 | -0,25558435 | 1,51332437 | -0,690711305 | -0,157245427 | -1,219474462 |
| GSSG | 1,7103289 | -0,0992663 | 0,57798934 | -0,530912363 | -1,021771904 | -0,636367671 |
| GTP | 0,63212332 | 0,25123902 | -0,62343379 | -0,898080463 | -0,933613532 | 1,571765448 |
| guanine | 0,80027108 | 0,7394882 | -1,73590314 | 0,575677356 | -0,634702953 | 0,255169454 |
| Hexanoic acid | 0,46665627 | -1,19416835 | -0,56722248 | -0,734611426 | 0,547941778 | 1,481404202 |
| Hexanoyl-carnitine | 0,85881106 | 0,7426243 | 1,094243 | -1,042435911 | -1,000268911 | -0,65297354 |
| Histidine | -0,61812893 | -1,0301213 | 0,35520931 | 1,24323238 | 0,995400506 | -0,945591968 |
| Hydroxy-L-proline | -1,01124801 | -0,78284162 | -0,92558014 | 0,943120707 | 1,030221555 | 0,746327507 |
| hypoxanthine | -0,32517573 | -0,53989046 | -1,24837401 | 0,334685984 | 0,0676888 | 1,711065422 |
| IMP | 0,72920143 | 1,24208441 | 0,71715475 | -0,921454838 | -0,861330496 | -0,90565525 |
| Inosine | 0,88494425 | -0,35516317 | -1,1109352 | -0,18837936 | -0,738838641 | 1,50837212 |
| IsoLeucine | -1,67894498 | 0,38552024 | 1,36073973 | -0,100228387 | 0,308276764 | -0,275363368 |
| Lactate | -1,39971421 | -0,27641615 | 0,61860899 | -0,857589361 | 0,878710308 | 1,036400422 |
| L-Alanine | -1,09557574 | -1,1646498 | 1,25278145 | 0,93060342 | 0,087757492 | -0,010916823 |
| L-alpha-Amino-N-butyric aci | 0,53175312 | 0,19039589 | 0,31422843 | -0,973172579 | -1,379412917 | 1,316208058 |
| L-beta-Aminoisobutyric acid | 0,4247825 | -0,19751142 | -0,58682916 | 0,108875791 | -1,356702353 | 1,607384643 |
| L-Dihydroorotic acid | -0,9191962 | -0,46631872 | -0,20314524 | -0,072564676 | 1,951160854 | -0,28993601 |
| Leucine | -0,42153554 | 0,01041418 | -0,06036559 | -0,785824316 | -0,677219487 | 1,934530754 |
| Linoleic acid | 0,10106294 | 0,83055379 | 0,59484551 | -1,815154633 | 0,696463718 | -0,407771325 |
| L-Kynurenine | 0,16011858 | -0,84687369 | -0,76289476 | -0,26921697 | -0,171447077 | 1,890313918 |
| L-Sarcosine | -0,31066565 | -1,25369273 | -0,93451832 | 1,083039775 | 1,084179433 | 0,331657495 |
| Lysine | 1,26257175 | 0,85875971 | 0,51864012 | -0,671402363 | -0,909976904 | -1,058592306 |
| Malate | 0,33845774 | 1,40874395 | -0,49761788 | 0,247409196 | 0,109279101 | -1,6062721 |
| Methionine | -0,66379336 | -1,64415754 | 0,96276335 | 0,440616942 | 0,048587514 | 0,855983099 |
| methyl-lysine (N) | 0,32384528 | 0,13251019 | 1,776186 | -0,617001748 | -0,943882192 | -0,67165753 |
| Myristic acid | 0,50289074 | 1,03557281 | 0,38250368 | -1,822369778 | 0,268890261 | -0,367487712 |
| Myristoyl-carnitine | 0,34554609 | 0,67613406 | 1,23025599 | -1,180657979 | 0,150705987 | -1,221984152 |
| NAD+ | 1,12894861 | -0,21715794 | -1,26990539 | 0,015781899 | 1,172868165 | -0,830535343 |
| NADH | -0,36745857 | -0,42919353 | -0,39991473 | -0,422974378 | -0,421178456 | 2,040719656 |
| NADP+ | 0,89974473 | 1,22303724 | -0,8682871 | -1,239072457 | 0,442453965 | -0,457876373 |
| NADPH | 1,32659134 | 0,3792099 | -0,64076598 | -1,28305284 | -0,603656563 | 0,821674138 |
| N-carbamoyl-L-aspartic acid | 0,46384911 | -1,82007062 | 1,15178438 | -0,2206082 | 0,269880939 | 0,155164397 |
| nicotinamide | -0,49040298 | 0,43628577 | 1,73035976 | -0,252709237 | -1,210730192 | -0,212803128 |
| nicotinamide N-oxide | -0,46601986 | 0,46079012 | -1,08748508 | 0,056866544 | 1,711270173 | -0,675421895 |
| Octanoic acid | 0,48118986 | -0,49897142 | 1,81106454 | -0,468454817 | -0,39549243 | -0,929335729 |
| Octanoyl-carnitine | 0,57584408 | 1,26967216 | 0,64366269 | -1,308704435 | -0,93124108 | -0,249233418 |
| Oleic acid | 1,01283964 | 0,56104366 | 1,00544245 | -1,09086395 | -0,325466084 | -1,162995713 |
| O-Phosphoethanolamine | 1,00288162 | 1,02875772 | 0,67819426 | -0,781212791 | -1,018421483 | -0,910199326 |
| Ornithine | 0,36070372 | 1,27498583 | 0,31853145 | 0,252981436 | -0,536604399 | -1,670598034 |
| Orotic acid | -0,47166414 | 1,58351878 | -0,61870763 | -0,322635385 | 0,855108848 | -1,025620471 |
| oxypurinol | 0,43941305 | 0,7534663 | -1,44872624 | -0,039297074 | -0,875975219 | 1,171119184 |
| Palmitic acid | 1,21749029 | 0,97947363 | 0,12142159 | -1,476897531 | -0,356213896 | -0,485274085 |
| Palmitoleic acid | 0,97910878 | 0,44577688 | 1,1869253 | -1,182903049 | -0,797184685 | -0,631723234 |
| Palmitoyl-carnitine | 0,22278552 | 0,91421686 | 1,38613969 | -0,738899397 | -0,72560162 | -1,058641052 |
| Pantothenate | 0,04413153 | 0,8388821 | 1,43172953 | -0,319418412 | -1,273153967 | -0,72217078 |
| PEP | 0,67330151 | 0,7036675 | 1,29159443 | -0,813018732 | -0,908043722 | -0,947500979 |
| Phenylalanine | 0,95636851 | 0,47217876 | 1,2260063 | -0,961900021 | -0,873943728 | -0,818709818 |
| phosphocreatine | 0,83414643 | 0,13602025 | -1,62666996 | -0,72491006 | 0,407866568 | 0,973546776 |
| Proline | 0,40239654 | -0,70973458 | -1,33975819 | 0,569868769 | -0,36538902 | 1,442616476 |
| Propionyl-carnitine | 0,89729034 | 0,63184128 | 1,16622147 | -0,890107665 | -0,985769869 | -0,81947557 |
| pyridoxal | -1,69446024 | -0,12056177 | 0,65945553 | 1,146168665 | 0,43227405 | -0,422876226 |
| Pyruvate | 0,12834773 | 0,23624802 | 1,81873441 | -0,852539096 | -0,603777562 | -0,727013498 |
| Quinolinic acid | -1,1246873 | 0,88255677 | 0,80314174 | -1,130279891 | -0,375505543 | 0,944774223 |
| riboflavin | -0,69043917 | 0,17109772 | -0,6991889 | -1,114252574 | 1,020348171 | 1,312434762 |
| Ribose phosphate | 0,47558996 | 1,06461971 | 1,10520221 | -1,092162122 | -0,876165388 | -0,677084372 |
| Sedoheptulose-7-phosphate | -0,47957083 | -0,01527055 | 0,33809584 | 1,329761909 | 0,462114578 | -1,635130946 |
| Serine | -0,11481759 | -0,72930437 | 1,87214464 | 0,249719206 | -0,450012967 | -0,827728917 |
| Serotonin | 0,54242817 | 0,85635451 | 0,22722815 | -0,118992744 | -1,93059973 | 0,423581637 |

| | | | | | | |
|---|---|---|---|---|---|---|
| Stearic acid | 1,1877037 | 0,99924691 | 0,18965053 | -1,376671769 | -0,782580749 | -0,217348617 |
| Succinic acid | -0,06698798 | 0,26111757 | 1,80384434 | -0,199960238 | -0,806635023 | -0,991378668 |
| Taurine | 0,81308495 | -0,94580317 | -0,26907534 | 0,272580293 | 1,34704564 | -1,217832385 |
| Threonine | 0,12959411 | -0,1400673 | 1,90803811 | -0,429875108 | -0,559019596 | -0,90867022 |
| Tryptophan | 1,18434814 | 0,35694916 | 1,07194895 | -0,757926014 | -1,040074109 | -0,815246123 |
| Tyrosine | 1,6166027 | -0,72954451 | 0,8857395 | -0,540590432 | -0,712074123 | -0,520133132 |
| UDP | 1,45304753 | 0,64360105 | -0,27147255 | -1,469951555 | -0,475003761 | 0,119779292 |
| UDP-GlcNAc | -0,27311011 | 0,10425332 | 1,35372882 | -1,501900978 | 0,781411006 | -0,464382057 |
| UMP | 0,18827679 | 0,81884718 | 1,44395032 | -0,976757167 | -1,027148723 | -0,447168393 |
| urate | 1,12652189 | 1,24159529 | -1,35248293 | -0,238433402 | -0,363023077 | -0,41417778 |
| uridine diphosphate galactos | 0,79042803 | 1,27612177 | 0,24071652 | -1,43461518 | -0,789816455 | -0,082834674 |
| UTP | 1,07471947 | 0,43275878 | -0,64117222 | -1,272234108 | -0,676134264 | 1,082062339 |
| Valine | 0,74123008 | 1,16873246 | 0,79875586 | -0,879325631 | -0,908413066 | -0,920979703 |
| Xanthine | -0,13828182 | 0,12077178 | -1,84748193 | 0,322604384 | 0,410951168 | 1,13143642 |
